# Nora virus proliferates in dividing intestinal stem cells and thereby sensitizes *Drosophila* flies to *Pseudomonas aeruginosa* intestinal infection and to oxidative stress

**DOI:** 10.1101/2025.01.30.635658

**Authors:** Adrien Franchet, Samantha Haller, Miriam Yamba, Vincent Barbier, Angelica Thomaz-Vieira, Vincent Leclerc, Stefanie Becker, Kwang-Zin Lee, Igor Orlov, Danièle Spehner, Laurent Daeffler, Dominique Ferrandon

**Affiliations:** UPR 9022 CNRS, IBMC, University of Strasbourg, France; UMR 7104 CNRS, U964 INSERM, IGBMC, University of Strasbourg, France; Institute for Parasitology and Research Center for Emerging Infections and Zoonoses, University of Veterinary Medicine Hannover, Hannover, Germany; Université Bourgogne Europe, Institut Agro, CNRS, INRAE, UMR CSGA, 21000 Dijon, France; Fraunhofer Institute for Molecular Biology and Applied Ecology (IME), Ohlebergsweg 12, Giessen, Germany; UMR 7178 CNRS, Institut Pluridisciplinaire Hubert Curien, Strasbourg, France; Institute of Translational Medicine and Liver Disease, Inserm U1110, Strasbourg, France

## Abstract

The digestive tract represents the most complex interface of an organism with its biotope. Food may be contaminated by pathogens and toxicants while an abundant and complex microbiota thrives in the gut lumen. The organism must defend itself against potentially noxious biotic or abiotic stresses while preserving its microbiota, provided it plays a beneficial role. The presence of intestinal viruses adds another layer of complexity. Starting from a differential sensitivity of two lines from the same *Drosophila* wild-type strain to ingested *Pseudomonas aeruginosa*, we report here that the presence of Nora virus in the gut epithelium promotes the sensitivity to this bacterial pathogen as well as to an ingested oxidizing xenobiotic. The genotype, age, nature of the ingested food and, to a limited extent, the microbiota are relevant parameters that influence the effects of Nora virus on host fitness. Mechanistically, we detect the initial presence of the virus essentially in progenitor cells. Upon stress such as infection, exposure to xenobiotics, aging or feeding on a rich-food diet, the virus is then detected in enterocytes, which correlates with a disruption of the intestinal barrier function in aged flies. Finally, we show that the virus proliferates only when ISCs are induced to divide. We propose that enterocytes essentially get infected through lineage from progenitor cells and are not directly infected.

In conclusion, it is important to check that experimental strains are devoid of intestinal viruses when monitoring survival/life span of fly lines or when investigating the homeostasis of the intestinal epithelium as these viruses can constitute significant confounding factors.

## Introduction

The digestive tract represents the most complex interface of the host with its environment as it involves a large surface, the presence of an important microbiota, the ingestion of food potentially contaminated by pathogens or toxicants, and the need for absorption of digested nutrients and solutes. Multiple concurrent infections or abiotic stresses are likely to be common, especially when the host feeds on decaying organic matter, as is the case for *Drosophila melanogaster*. Co-infection models with bacterial and viral pathogens in the gut are starting to be experimentally investigated and have revealed an enhanced susceptibility to combined infections that can be caused either by affecting pathogen transmission or by an inability to withstand tissue damage (Lian et al., 2022, and references therein).

Some twenty distinct viruses have been identified in wild Drosophila and around 30% of wild individuals are infected by at least one virus (Webster et al., 2015). A few of them are studied in the laboratory for the mechanisms of interaction with the host: *Drosophila melanogaster* Sigma Virus (DMelSV), Drosophila A virus (DAV), Drosophila C virus (DCV), Drosophila X virus (DXV), and Nora virus (Schneider and Imler, 2021; Webster et al., 2015). Nora is an enteric virus (Habayeb et al., 2006). Unlike other insect picorna-like viruses, its genome encodes four open reading frames corresponding to: a suppressor of RNA interference, VP1 (van Mierlo et al., 2012), replicative proteins coded by ORF2, the poorly characterized product of ORF3, and capsid proteins derived from ORF4 (Ekstrom et al., 2011). This virus infects common laboratory stocks where it appears to cause a persistent infection. It is transmitted horizontally and vertically via a fecal-oral route (Habayeb et al., 2009a; Habayeb et al., 2006). The Nora virus likely proliferates in the digestive tract, as large quantities of the virus are continuously released in the feces of infected flies. The virus does not appear to have major effects on host fitness, even though some damages to the intestinal epithelium have been reported (Habayeb et al., 2009a). However, the microbiota of flies infected by Nora may change in quantity and bacterial diversity (Schissel et al., 2021). We then wondered whether a persistent Nora virus infection could influence a secondary pathogenic bacterial infection.

The *Drosophila* intestinal epithelium is simple, formed mostly by a monolayer of columnar polyploid enterocytes (EC) and also comprises enteroendocrine cells and intestinal stem cells (ISCs) and enterocytes progenitor called enteroblasts (Jiang and Edgar, 2011; Lemaitre and Miguel-Aliaga, 2013). The microbiota is made up of few species, at most twenty, and is usually dominated by two-three prevalent species such as *Acetobacter pomorum*, *Lactobacillus plantarum*, or *Enterococcus faecalis,* but its composition changes with ageing (Broderick and Lemaitre, 2012).

Immune defenses in the midgut include a chemical armamentarium, notably reactive oxygen species (ROS) generated by the NADPH oxidase, NOX, and likely not Dual oxidase, and antimicrobial peptides such as Diptericin (Ayyaz and Jasper, 2013; Buchon et al., 2013; Ferrandon, 2013; Iatsenko et al., 2018; Jones et al., 2013; Kim and Lee, 2014; Liu et al., 2024; Patel et al., 2019). Resilience, a complementary dimension of mucosal host defense also known as disease tolerance (Medzhitov et al., 2012), is the ability of the intestinal epithelium to maintain its homeostasis, for instance by a mechanism of cytoplasmic purge (Lee et al., 2016) or the increased proliferation of ISCs that ultimately regenerate enterocytes damaged either directly by pathogens or indirectly by the host’s own immune response (Ferrandon, 2013, and references therein). ISC proliferation is regulated by the production of the growth factor Unpaired 3 (Upd3) that activates the JAnus Kinase/Signal Transducers and Activators of Transcription (JAK/STAT) pathway in ISCs (Buchon et al., 2009b; Herrera and Bach, 2019). Several models of gut infection have been developed in *Drosophila* (Bonfini et al., 2016) and can be used to study coinfection with Nora virus. We have focused on *Serratia marcescens* and *Pseudomonas aeruginosa* intestinal infections (Chen et al., 2025; Chen et al., 2024; Cronin et al., 2009; Haller et al., 2018; Lee et al., 2016; Limmer et al., 2011; Nehme et al., 2007; Sina Rahme et al., 2022; Socha et al., 2023) and had not noticed major damages of *P. aeruginosa* oral infections on the integrity of the intestinal epithelium in contrast to another study (Apidianakis et al., 2009). *P. aeruginosa* bacteria are able to cross the intestinal barrier and to silently colonize inner tissues (Chen et al., 2024). They are initially found circulating in the hemolymph at low concentration. However, after a few days of continuous feeding on bacterial solution, the bacteria start proliferating in the hemolymph in a quorum-sensing-dependent manner and ultimately kill the host through bacteremia (Haller et al., 2018; Limmer et al., 2011).

Here, we demonstrate that Nora virus is a co-factor that synergizes with ingested bacteria or toxicants in *Drosophila* intestinal infection models, thus leading to an earlier demise of infected flies. We also report major effects of Nora infection on the lifespan of the flies under different feeding conditions. Mechanistically, we find that Nora virus initially infects ISCs and proliferates intensely upon ISCs divisions, ultimately leading to the contamination of enterocytes and gut barrier function disruption. Blocking the compensatory ISC proliferation by either inhibiting apoptosis in enterocytes or interfering with JAK-STAT signaling in ISCs protects flies from Nora pathogenesis through a drastic decrease of its load.

## Results

### Nora virus-infected stocks are shorter-lived and more susceptible to some infections

We noted that two Oregon-R (Ore-R) wild-type stocks kept by different investigators in the laboratory, hereafter referred to as (SM) and (SC), displayed a remarkably distinct survival pattern in one out of two models of intestinal infections (Limmer et al., 2011; Nehme et al., 2007) in which flies respectively continuously feed either on *Pseudomonas aeruginosa* PA14 (Fig. 1A) or *Serratia marcescens* Db11 (Fig. S1A). Namely, the Ore-R (SM) was more sensitive to PA14 and there was a slight trend toward higher susceptibility to Db11. We also noted that Ore-R (SM) displayed a shorter lifespan under non-infected conditions either on our standard food or on sucrose solution (Fig. 1B, Fig. S1B, and see below). We wondered whether the difference might be due to an infection and checked the stocks for the presence of common microbes known to affect *Drosophila* stocks (Haller et al., 2014; Lestradet et al., 2014). The only notable difference we identified between the two stocks was the presence of the Nora enteric virus in the infection-sensitive Ore-R(SM) (Fig. 1C).

**Figure 1.**
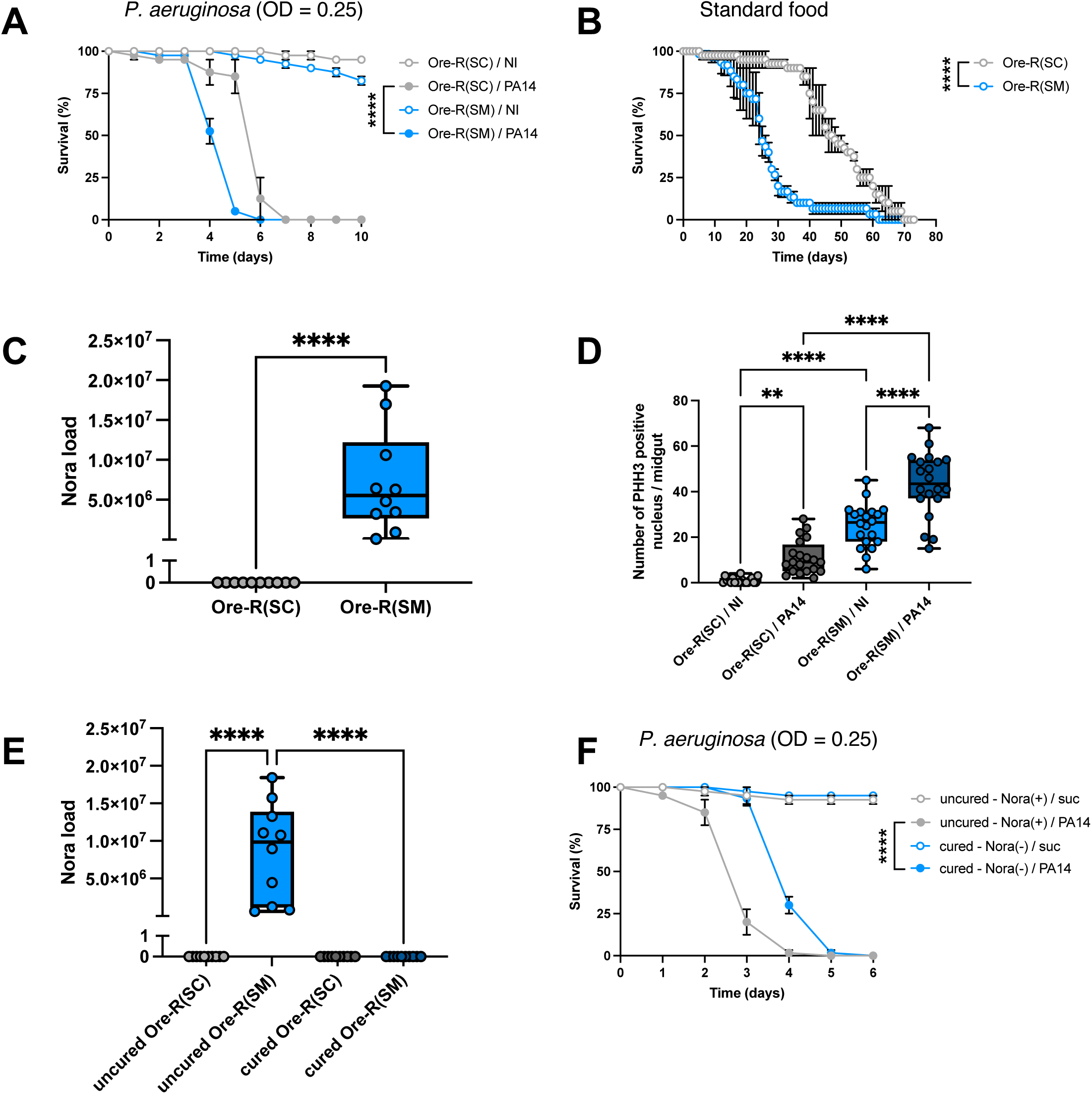
Correlation between Nora virus infection and impaired defense against *P. aeruginosa* intestinal infection as well as lowered fitness. The Oregon-R sub-strain infected with Nora virus is referred to as Ore-R(SM), while the non-infected sub-strain is referred to as Ore-R(SC). (**A**) Survival of Ore-R(SM) (Nora(+)) and Ore-R(SC) (Nora(-)) flies following oral exposure to *P. aeruginosa* PA14 (OD 0.25) at 25 °C. NI: not infected. (**B**) Lifespan analysis of Nora-infected and non-infected flies maintained under standard laboratory conditions at 25 °C. (**C**) Nora virus RNA levels in Ore-R(SM) and Ore-R(SC) stocks quantified by RT-qPCR. (**D**) Quantification of phospho-histone H3 (PHH3)-positive nuclei in the midgut of Ore-R(SM) (Nora(+)) and Ore-R(SC) (Nora(-)) flies at 2 days following oral PA14 infection or in non-infected (NI) controls. PHH3-positive nuclei counts were assessed as a measure of epithelial cell proliferation. (**E**) Nora virus RNA levels measured by RT-qPCR in the original Ore-R(SM) stock and in stocks cured of Nora virus by embryo bleaching. (**F**) Survival of uncured and cured Ore-R(SM) stocks to the ingestion of PA14. Each graphic is representative of three independent experiments. Survival panels are presented as mean ± SEM with each curve representing a triplicate of 20 flies. Other panels are presented as box and whiskers where the middle bar of the box plots represents the median, and the upper and lower limits of boxes indicate the first and third quartiles, respectively; the whiskers define the minima and maxima. Each dot in Nora virus load panels (C, E) represent a sample of 5 flies. Each dot in the PHH3 quantification panel (D) represents one single posterior midgut. Survival data were analyzed using a log-rank (Mantel-Cox) test. Nora virus load in (C) was analyzed using a Mann-Whitney nonparametric test. PHH3 quantification (D) and Nora load (E) were analyzed using one-way ANOVA with Tukey’s post-hoc tests. Statistical significance is indicated as **p < 0.01, ***p < 0.001, ****p < 0.0001.

We therefore examined the guts of flies from the two Ore-R stocks and did not detect any major morphological differences. However, when we measured basal ISC proliferation by counting phospho-histone H3-positive (PHH3) cells in the intestine, we found an enhanced rate of ISC proliferation in the Nora-positive Ore-R(SM) stock (Fig. 1D). When we challenged the Nora-negative Ore-R(SC) stock with *P. aeruginosa*, we found a small but nevertheless significant increase in ISC proliferation. However, this *P. aeruginosa*-induced increase was three-times larger in the Nora-positive Ore-R(SM) stock (Fig. 1D).

Overall, these observations suggest that flies infected with Nora have a lower fitness and are more susceptible to infections than non-infected flies.

### Nora virus causes the enhanced susceptibility to *P. aeruginosa* intestinal infections

As expected, bleaching the eggs laid by the Nora-infected Ore-R(SM) stock appeared to eradicate the virus ((Habayeb et al., 2009b), Fig. 1E and Fig. S2A). This treatment did not have a noticeable adverse impact as bleached stocks displayed a normal survival for at least ten days on sucrose (Fig. S1E-I). Furthermore, cured Ore-R(SM) flies fed with *P. aeruginosa* were less susceptible to this intestinal challenge than the uncured stock (Fig. 1F).

We placed our cured flies in a vial that had hosted infected males for several days for fecal-oral transmission (Habayeb et al., 2009a) and found that they became again Nora-positive over several generations (Fig. S2A). The reinfected stock was more sensitive to the ingestion of *P. aeruginosa* than the cured stock (Fig. S2B). This correlated with an enhanced ISC proliferation in the Nora virus reinfected flies, whether challenged with *P. aeruginosa* or not (Fig. S2C).

The drawback of the fecal transmission route is that other enteric pathogens may be transferred along with the Nora virus. To exclude this possibility, we used gradient centrifugation to purify and concentrate Nora virus from an infected fly extract. This preparation was homogenous with particles of the expected size when observed by cryo-electron microscopy (Fig. 2A). Using RT-PCR, we confirmed the identity of the virus and ruled out the presence of other *Drosophila* viruses (DAV, DCV, DBV, DTV, DXV, CrPV, FHV…; see Table 1 Primer Sequence). Cured flies were fed on this pure viral preparation for 24 hours. This was sufficient to stably reinfect the stock over several generations (Fig. 2B). As with the fecal contamination route, we observed that the flies reinfected with the pure viral preparation were more sensitive to the ingestion of *P. aeruginosa* (Fig. 2C) and displayed an enhanced rate of ISC proliferation in the gut (Fig. 2D). In terms of lifespan on standard food or on sucrose solution, the re-infected stocks displayed a largely decreased fitness (Fig. 2E-F), in contrast to the findings of (Habayeb et al., 2009a; Nigg et al., 2024) who reported only a mild effect upon the injection of the Nora virus. Altogether, these results demonstrate that Nora virus is responsible for the enhanced sensitivity to *P. aeruginosa* intestinal infection.

**Figure 2.**
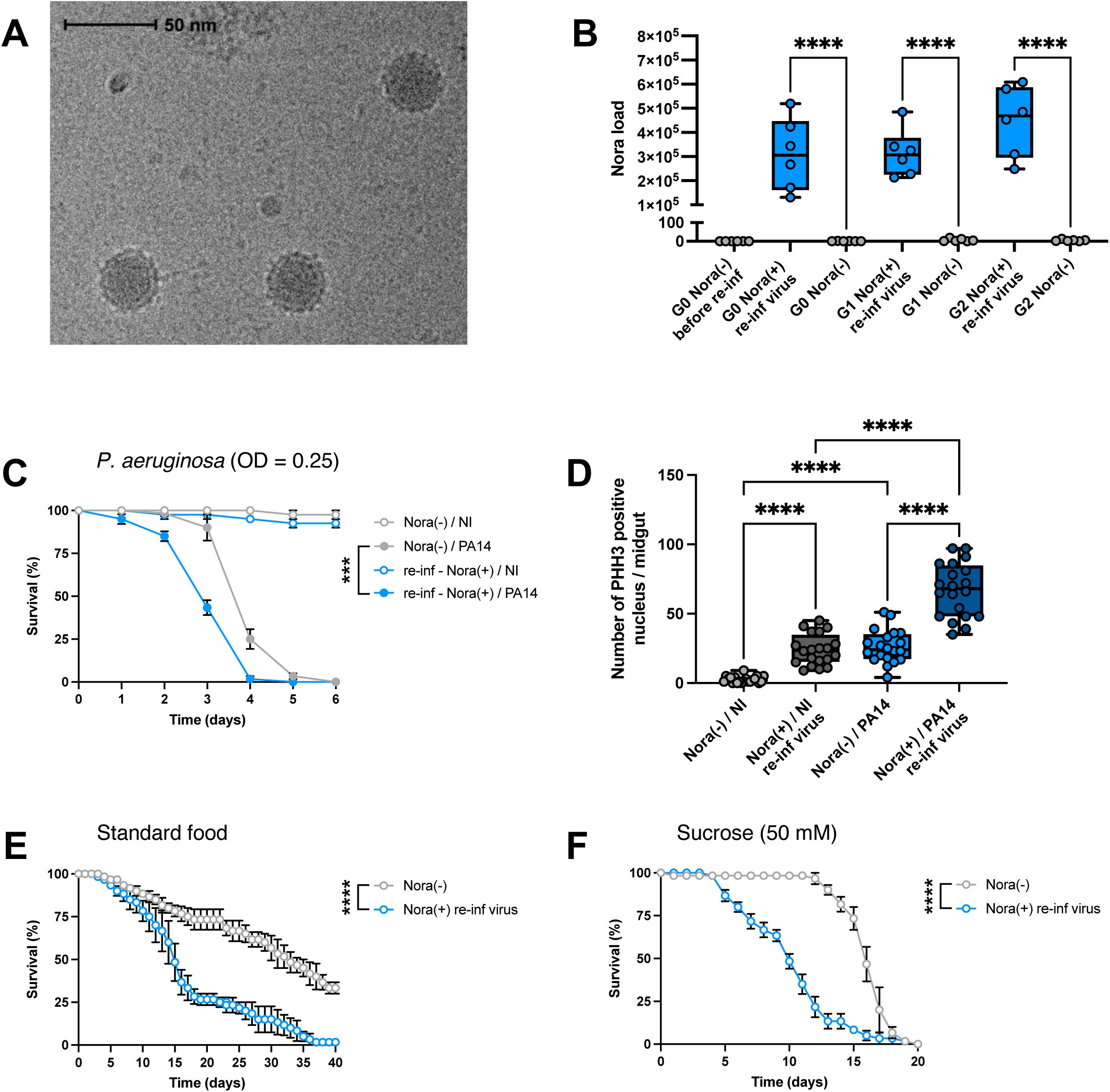
Infection of Nora(-) flies with the purified Nora virus recapitulates the fitness properties of Nora naturally infected flies. Flies used in these experiments were derived from Ore-R(SM) stocks cured of Nora virus by egg bleaching. A subset of cured flies was experimentally re-infected with a purified Nora virus preparation (Nora(+) re-inf virus), while control flies remained uninfected (Nora(-)). PA14: infection with *P. aeruginosa* PA14; NI: noninfected. (**A**) Transmission electron microscopy image of the purified Nora virus preparation used for re-infection. (**B**) Nora virus RNA levels measured by RT-qPCR in Nora(+) and Nora(-) flies across successive generations following experimental re-infection: G0 to G2. (**C**) Survival of re-infected Nora(+) and Nora(-) flies following oral exposure to PA14 at 25 °C. (**D**) Quantification of phospho-histone H3 (PHH3)-positive nuclei in the midgut of re-infected Nora(+) and Nora(-) flies following two days of PA14 ingestion or in non-infected (NI) controls. (**E**) Lifespan analysis of Nora(+) and Nora(-) flies maintained under standard laboratory conditions at 29 °C. (**F**) Lifespan analysis of re-infected Nora(+) and Nora(-) flies maintained on a sucrose-only diet at 25 °C. Each graphic is representative of three independent experiments. Survival panels are presented as mean ± SEM with each curve representing a triplicate of 20 flies. Other panels are presented as box and whiskers where the middle bar of the box plots represents the median, and the upper and lower limits of boxes indicate the first and third quartiles, respectively; the whiskers define the minima and maxima. Each dot in the Nora virus load panel (B) represents a sample of 5 flies. Each dot in the PHH3 quantification panel (D) represents one single posterior midgut. Survival data were analyzed using a log-rank (Mantel-Cox) test. Nora virus load was analyzed using a Mann-Whitney nonparametric test (B). PHH3 quantification (D) was analyzed using one-way ANOVA with Tukey’s post-hoc test. Statistical significance is indicated as ***p < 0.001, ****p < 0.0001.

We suspected that a polymorphism in the gene *pastrel* might be the cause of the different susceptibility to Nora virus between Ore-R(SM) and Ore-R(SC) (Magwire et al., 2012). To test this hypothesis, we determined by RT-PCR the presence of a *pastrel* sensitive or resistant alleles (*pst^S^*, *pst^R^*) in several stocks commonly used in the laboratory. Strikingly, the sensitive allele was found in homozygous condition only in the Ore-R(SM) stock, while it was heterozygous in our *w* (A5001) stock (Fig. S1D). Interestingly, the *w* (A5001) (Thibault et al., 2004) and the DD1 *cn bw* (Jung et al., 2001) stocks were found to be also harboring the Nora virus, the latter one being *pst^R^/ pst^R^* (Fig. S1C-D). The *w* (A5001) and DD1 *cn bw* cured stocks as well as the Canton-S stocks were all more susceptible to *P. aeruginosa* ingestion after having been converted to a Nora-positive status by the prior ingestion of the purified virus preparation (Fig. S1E-G). Since even stocks harboring two copies of the resistant *pastrel* allele became more sensitive to ingested *P. aeruginosa* (Fig. S1D, S1F-G), we can rule out that distinct *pastrel* alleles were also the cause of the enhanced sensitivity to bacterial intestinal infections. Unexpectedly, the Ore-R(SC) strain appeared to be resistant to the ingestion of Nora since it did not display an enhanced sensitivity to oral *P. aeruginosa* infection (Fig.S1I). Thus, this line may harbor in addition to the homozygous *pst^R^* allele a polymorphism at an unidentified locus that restricts the Nora virus.

In the following, we will use the Ore-R(SM) cured stock as a Nora-negative control. We will further characterize the impact of Nora virus on infected flies using the cured Ore-R(SM) stock reinfected with the pure Nora preparation, which will be referred to as Nora-positive flies.

### Nora virus is reducing host longevity largely independently of the microbiota

We confirmed that Nora re-infected flies succumbed much earlier than Nora-negative control flies when orally challenged with *P. aeruginosa* in a sucrose solution after five days of prior feeding on our standard food or a rich food corresponding to our standard food containing five times more yeast (Fig. S3A-B). Strikingly, we observed that an oral infection with PA14 increased the Nora burden (thrice on standard food and four-times on rich food). Actually, both the Nora virus load and the *P. aeruginosa* titer was higher for flies fed on rich food (Fig. 3A, Fig. S3C). Of note, flies fed on rich food succumbed earlier to the challenge, independently of their Nora status (compare S3B to S3A). We noted a correlation between the Nora load in the absence or presence of PA14 and the measured proliferation of ISCs, which held also on rich food (compare Fig. 3A and Fig. S3D).

**Figure 3.**
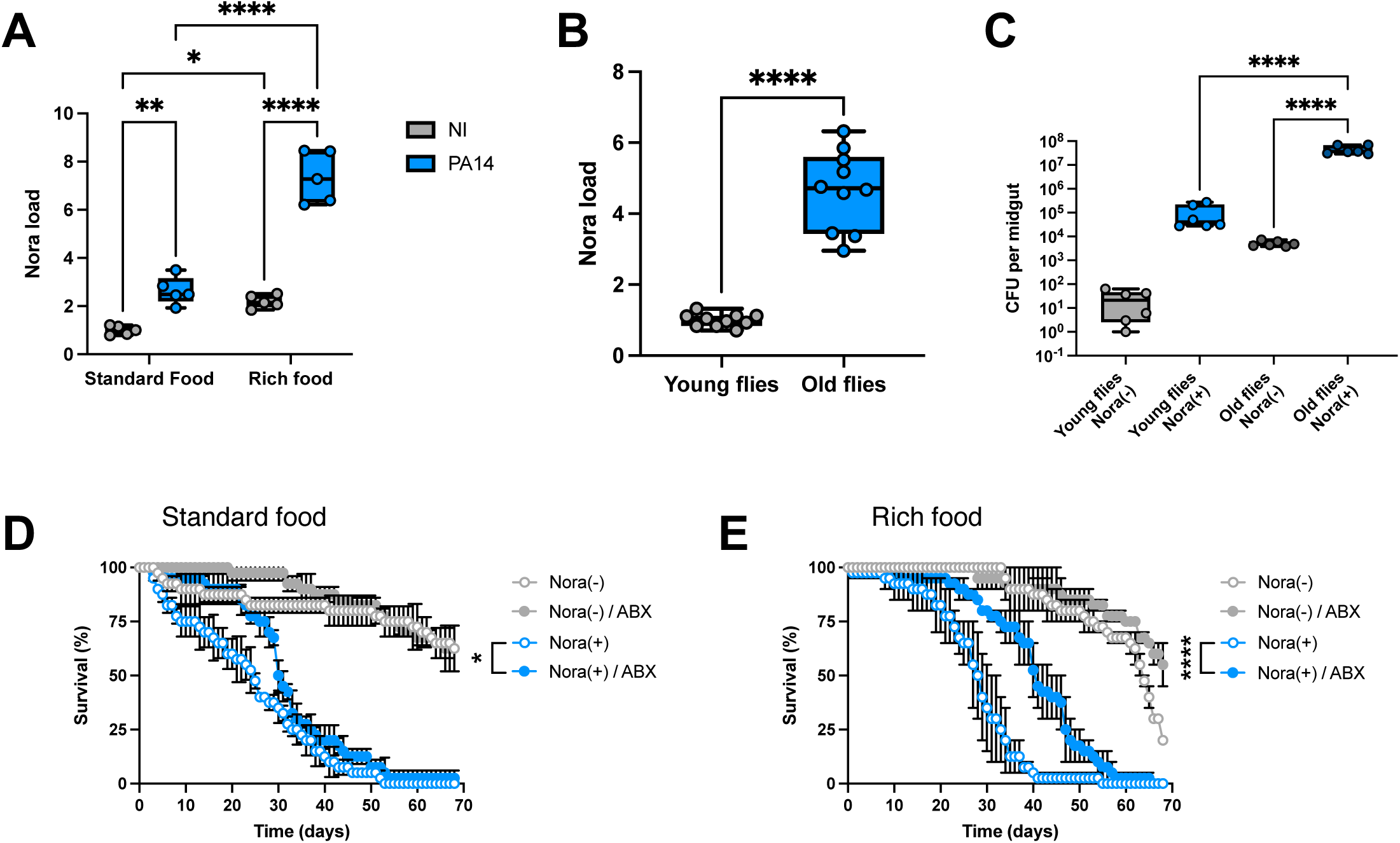
Influence of age, diet, and microbiota on Nora virus load and fly fitness. (**A**) Nora virus RNA levels measured by RT-qPCR at 3 days following oral exposure to *P. aeruginosa* PA14 in Nora-infected and non-infected flies maintained on either standard diet or rich diet (standard medium supplemented with 5x yeast). (**B**) Nora virus RNA levels measured by RT-qPCR in young (3-day-old) and aged (30-day-old) Nora-infected flies. (**C**) Quantification of total culturable bacterial microbiota by colony-forming unit (CFU) counts in young Nora(+) and Nora(-) flies maintained for 8 days on a sucrose-only diet, and in aged flies maintained on standard fly food. CFU values are presented on a logarithmic scale. (**D**) Lifespan analysis of Nora-infected (Nora(+) and non-infected (Nora(-) flies maintained on standard diet in the presence or absence of antibiotics (ABX). (**E**) Lifespan analysis of Nora-infected (+) and non-infected (-) flies maintained on rich diet in the presence or absence of antibiotics (ABX). Each graphic is representative of three independent experiments. Survival panels are presented as mean ± SEM with each curve representing a triplicate of 20 flies. Other panels are presented as box and whiskers where the middle bar of the box plots represents the median, and the upper and lower limits of boxes indicate the first and third quartiles, respectively; the whiskers define the minima and maxima. Relative unit was used for Nora virus load panels. Each dot in Nora virus load panels (A, B) represent a sample of 5 flies. Each dot in the CFU quantification panel (C) represents one single posterior midgut microbiota. Survival data were analyzed using a log-rank (Mantel-Cox) test. Nora virus load was analyzed using a Mann-Whitney nonparametric test (A, B). CFU quantification was analyzed using one-way ANOVA with Tukey’s post-hoc test (C). Statistical significance is indicated as *p < 0.05, **p < 0.01, ****p < 0.0001.

As aging flies exhibit a dysbiosis that correlates with an impaired homeostasis of the intestinal epithelium (Biteau et al., 2008; Buchon et al., 2009a; Choi et al., 2008), we also compared the Nora burden in young (3-5 day-old) and old (30-35 day-old) flies and found an about four-fold increase (Fig. 3B), which correlated with an enhanced rate of ISC proliferation (Fig. S3E), in keeping with a previously published study (Hanson and Lemaitre, 2023). While checking for an increase of the microbiota load in old flies (Iatsenko et al., 2018), we noted that the microbiota titer was much higher in Nora-positive than in Nora-negative flies (Fig. 3C). This observation also held when flies were kept feeding only on a sucrose solution (Fig. S3F). To determine whether the microbiota influenced the fitness of Nora-negative or -positive flies, we first monitored their survival when kept on sucrose solution. An antibiotic mix treatment substantially decreased the lifespan of Nora-negative flies (Fig. S3G), possibly reflecting a positive impact of the microbiota on this amino-acid depleted food solution even though the microbiota is minimal under these sucrose conditions (Fig. S3F). In contrast, antibiotics treatment had no effect on Nora-positive flies feeding on sucrose solution (Fig. S3G). Interestingly, the proliferation of ISCs was increased in antibiotics-treated Nora-positive flies kept under these amino-acid depleted conditions (Fig. S3H), which inversely correlates with the life span of those flies. We next monitored the fitness of Nora-negative or -positive flies on our standard or on rich food. The antibiotic treatment had a limited effect on Nora-negative flies with a somewhat enhanced fitness that failed to reach statistical significance (Fig. 3D-E). In contrast, it significantly extended the lifespan of Nora-positive flies on either standard or rich food (note that the detrimental effect of the Nora virus is attenuated on rich food, at least for about two-thirds of their lives, until they rapidly succumb). However, the antibiotics treatment failed by far to protect the flies to the level of Nora-negative flies. We conclude that under these different feeding conditions, the microbiota contributes at best only partially to the decreased fitness of Nora-positive flies.

### Nora virus contamination impairs the barrier function of the intestinal epithelium

As an increased rate of ISC division may mirror a phenomenon of compensatory proliferation driven by damage to enterocytes, we assessed the permeability of the gut of Nora-positive flies, first using the SMURF assay on 30-day old flies; the SMURF assay monitors the permeability of the gut barrier upon ingestion of a dye (Rera et al., 2011). The percentage of SMURF-positive flies was much higher in 30-day old Nora-positive flies fed on rich food (a time point corresponding to the death of 50% (LT50) of the flies in fitness experiments shown in Fig. 3E) (Fig. 4A-B). In keeping with this result, when we plated hemolymph collected from flies, we observed a strong microbial growth for that retrieved from SMURF-positive flies, but not from SMURF-negative flies, whether Nora-positive or Nora-negative (Fig. 4C). Thus, SMURF-positive flies do not only display an enhanced permeability to the dye but also likely to the midgut microbiota, an indication of severe disruption of the barrier function of the intestinal epithelium. These results were further confirmed after *P. aeruginosa* ingestion: already after three days, the passage of PA14 through the intestinal epithelium was much higher in Nora-positive than in Nora-negative flies (Fig. 4D). Of note, at Day 3, some 220 bacterial colonies on average were also retrieved from the hemolymph of Nora-positive control flies that had not ingested PA14, possibly reflecting an early intestinal barrier dysfunction in Nora-positive flies. Indeed, there was a detectable induction of a local immune response in the midgut induced by PA14 ingestion only in Nora-positive flies at two days instead of around five days for Nora-negative flies (Limmer et al., 2011), as monitored by measuring *Diptericin* expression levels (Fig. 4E). This immune response is triggered by the proliferation of *P. aeruginosa* in the hemolymph (Chen et al., 2024; Limmer et al., 2011). We conclude that the Nora virus affects the integrity of the intestinal epithelium and its barrier function.

**Figure 4.**
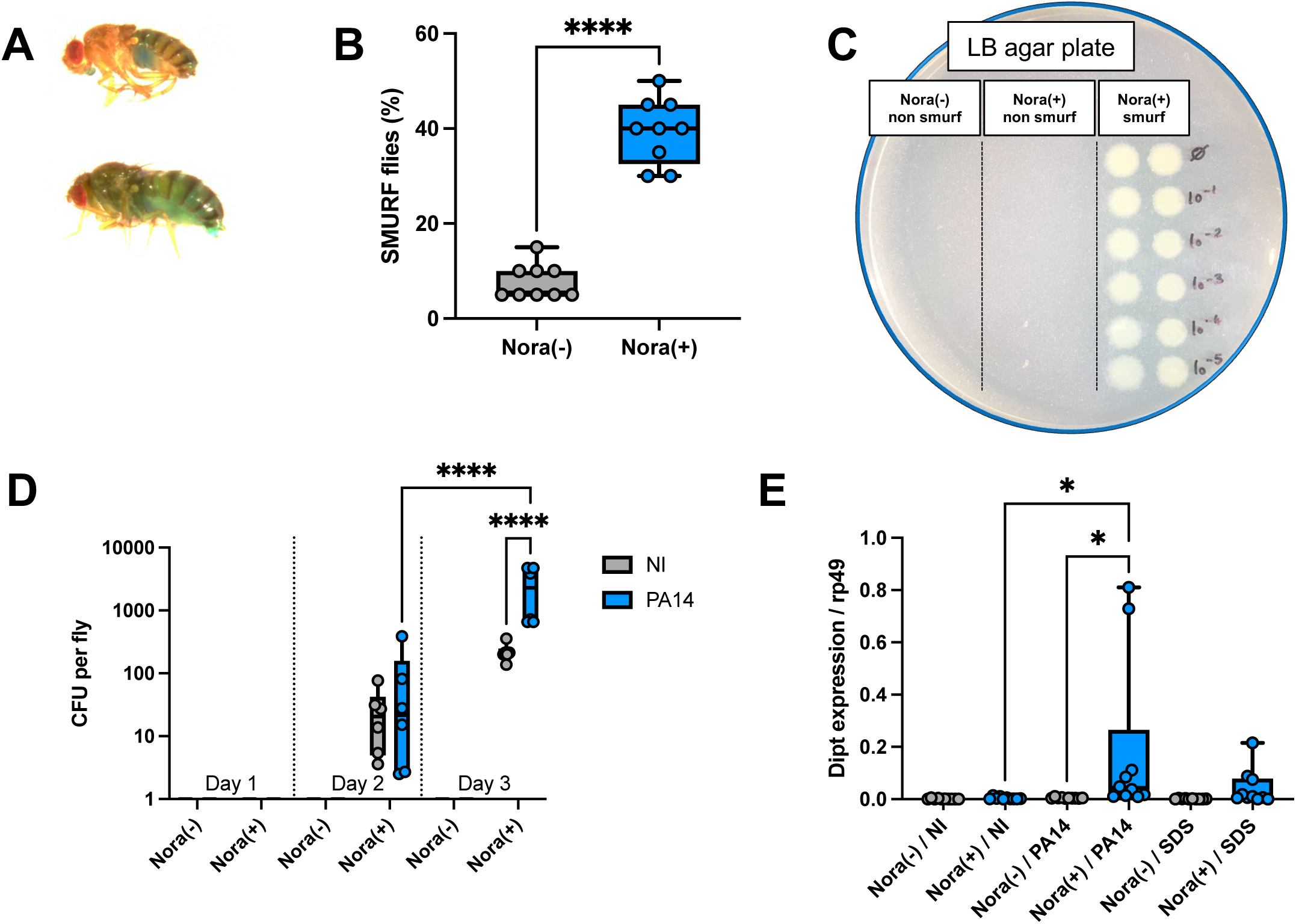
Alterations in intestinal barrier integrity and systemic bacterial dissemination in Nora virus-infected flies. (**A**) Representative images from the SMURF assay in 30-day-old non-infected fly (top) and Nora-infected fly (bottom) fed on a sucrose solution containing a blue dye. (**B**) Quantification of SMURF-positive Nora(-) or Nora (+) flies at 30 days of age maintained on rich diet (standard medium supplemented with 5x yeast) (**C**) Representative LB agar plate showing bacterial growth from serial dilutions of hemolymph collected from non-infected flies and from Nora-infected flies exhibiting either SMURF-negative or SMURF-positive phenotypes. Age of flies corresponding to the LT50 observed in the survival assay shown in Fig. 3E (Nora(-): 65 days; Nora(+): 30 days). (**D**) Quantification of bacterial titers in the hemolymph at 3 days following oral exposure to *P. aeruginosa* PA14 in Nora-infected and non-infected flies. (**E**) Relative expression of the antimicrobial peptide gene *Diptericin* in dissected midguts of Nora(+) flies measured by RT-qPCR following 2 days of PA14 intestinal infection or after feeding with 1% SDS for 4-6 hours. Each graphic is representative of three independent experiments. All panels are presented as box and whiskers where the middle bar of the box plots represents the median, and the upper and lower limits of boxes indicate the first and third quartiles, respectively; the whiskers define the minima and maxima. Each dot in the SMURF quantification panel (A) represents a single SMURF-positive fly. Each dot in the CFU quantification panel (C) represents one single fly hemolymph bacterial titer. Each dot in the *Diptericin* gene expression panel represents a sample of 5 midguts. The percentage of SMURF-positive flies was analyzed using a Mann-Whitney nonparametric test (B). CFU counts (D) and *Diptericin* gene expression levels (E) were analyzed using one-way ANOVA with Tukey’s post-hoc test. Statistical significance is indicated as *p < 0.05, ****p < 0.0001.

### Nora does not directly influence the survival of flies to septic injury

The experiments above show that the presence of the Nora virus in a stock increases its susceptibility to ingested *P. aeruginosa.* Since *P. aeruginosa* ultimately causes a systemic infection, we also tested whether Nora-positive flies would be more sensitive to other types of systemic infections. As shown in Fig. S4A, we did not observe any difference in the survival curves between Nora-positive or -negative flies exposed to spores of the entomopathogenic fungus *Beauveria bassiana*, an infection model in which no macroscopic wounds are inflicted. In contrast, Nora-positive flies succumbed significantly earlier than Nora-negative control flies when challenged with the Gram-positive *Enterococcus faecalis* or the Gram-negative *Enterobacter cloacae* bacterial pathogens in a septic injury model (Fig. S4B-C). However, we found in control experiments that Nora-positive flies pricked with a needle dipped into a sterile PBS solution succumbed at a rate that was similar to that observed in flies submitted to a septic injury, whereas control Nora-negative flies were more resistant to aseptic injury (compare Fig. S4D to Fig. S4B-C). Unexpectedly, *MyD88* flies displayed an enhanced apparent sensitivity to a clean injury, a phenomenon we have observed before and that depends on the endogenous microbiota of *MyD88* flies (Xu et al., 2023). To understand why Nora-infected flies were more susceptible to a near-sterile wound, we measured the proliferation rate of ISCs as it has been reported that injury triggers a ROS-response, in the hypodermis, hemocytes, and in the gut that leads to enterocyte apoptosis and an increased compensatory proliferation of ISCs (Chakrabarti and Visweswariah, 2020; Takeishi et al., 2013). As reported, we observed a modest, statistically not significant, increase of ISC proliferation in Nora-negative flies after injury of the cuticle (Fig. S4E). Nora-positive flies displayed a higher basal rate of ISC proliferation that was however not altered by injury, suggesting that gut damage is unlikely to account for the enhanced sensitivity of Nora-positive flies to wounding of the cuticle. We finally compared the survival rates of flies pricked with a clean needle to that of unchallenged controls and did not find any significant difference (Fig. S4F). We conclude that the apparent sensitivity of Nora-positive flies to clean injuries is actually due to the shortened lifespan of Nora-positive flies. This effect only becomes apparent in systemic infections with mild pathogens that kill the flies slowly (Fig. S4B-C).

### Nora virus is detected in intestinal stem cells in unchallenged flies and adopts an enterocyte localization upon challenge

Since the Nora virus is mostly detected in the gut by RT-qPCR (Habayeb et al., 2009a) and plating assays (Ekstrom and Hultmark, 2016) (Fig. 5A) and since it affects the barrier function of the intestinal epithelium, we raised an antibody against the virus, which is essentially specific except for a cross reaction of the secondary antibodies with intestinal muscles (compare Figs. 5B & S5A to 5C), which precludes drawing any conclusion as regards a possible localization of the Nora virus also in these muscles, as has been previously described for DCV oral infection (Ferreira et al., 2014). In contrast, a positive signal was detected in basally located small triangular cells of the intestinal epithelium of flies that had ingested the Nora virus five days earlier and not in non-infected controls (Figs. 5C-D, Fig. 5B & Fig. S5A). The right panel of Fig. 5C displays a dividing stem cell that yields a basally located ISC and a differentiating progenitor cell (enteroblast or enteroendocrine cell). Indeed, a similar non-basally located small cell is shown in Fig. S5B. We confirmed the localization of the Nora virus to progenitor cells of the intestinal epithelium by staining the Nora virus in *esg-Gal4Gal80^ts^ >UAS-GFP* flies (Fig. 5D). The Nora-positive signal was always found in the GFP-expressing progenitor cells. Using transgenic fly lines that express Dicer2-fluorescent protein fusions (Donelick et al., 2020; Girardi et al., 2015), we noted that even though the construct was expressed under the direct control of the poly-ubiquitin promoter (*ubi-p63E*), we failed to detect the fusion protein in ISCs (Figs. 5E & S5C), even though GFP is known to be stable in progenitor cells (Fig. 5D). If Dicer-2 were to be unstable specifically in ISCs, this might account for the initial localization of the Nora virus to ISCs, a proposition that would require further experimental confirmation.

**Figure 5.**
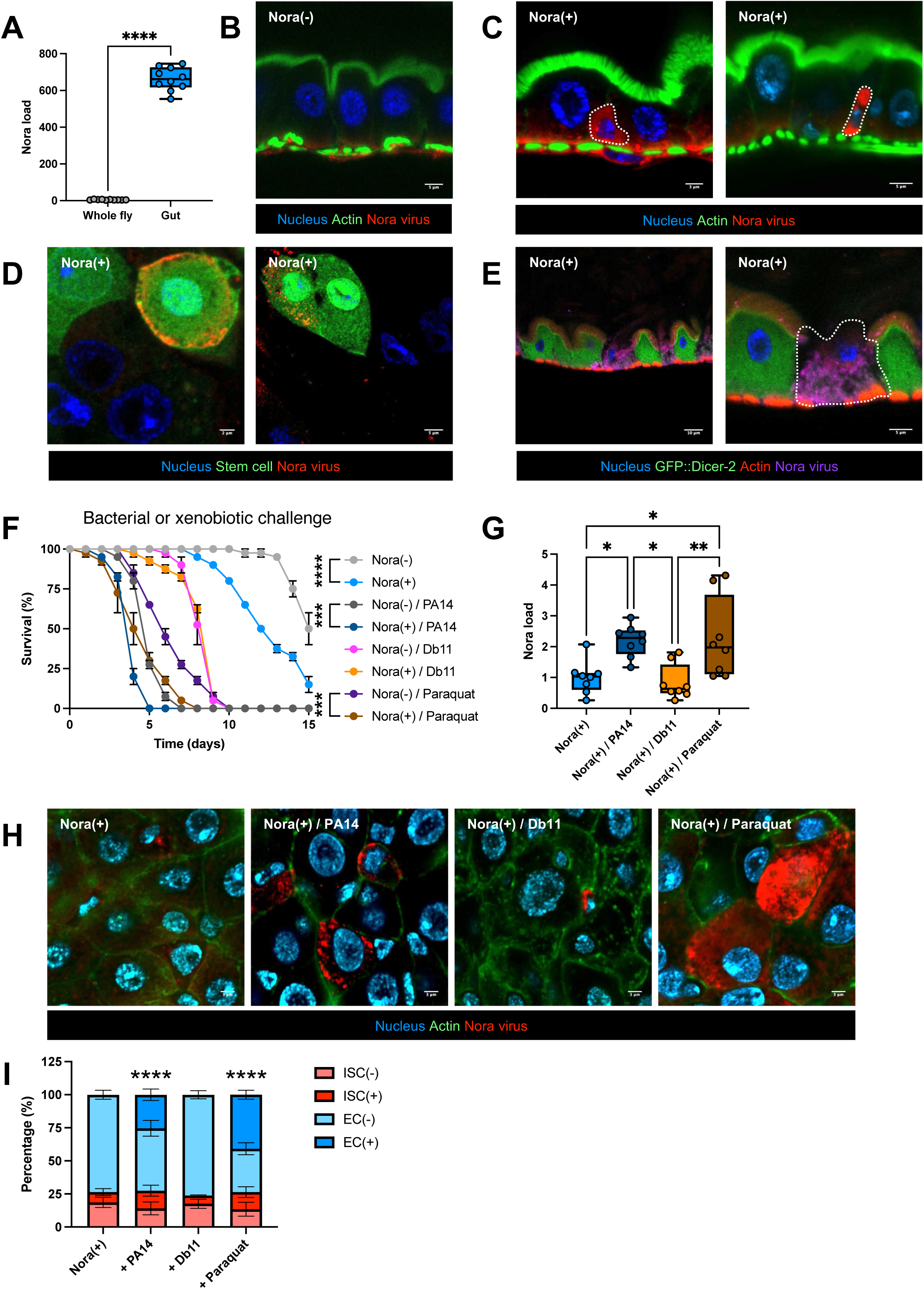
Nora virus is initially found in intestinal progenitor cells and is also located in enterocytes upon infectious or chemical stresses. (A) Nora virus RNA levels measured by RT-qPCR in whole flies or in dissected intestines. (B) Representative confocal images of intestines from 5-day-old non-infected flies. Tissues were fixed and stained for DNA (DAPI, blue), actin (FITC, green), and Nora virus (Cy3, red). The Cy3 secondary antibody (goat anti-mouse) shows background signal in intestinal muscles. Scale bar, 5 µm. (**C**) Representative confocal images of intestines from 5-day-old Nora-infected flies stained for DNA (DAPI, blue), actin (FITC, green), and Nora virus (Cy3, red). Scale bars, 3 µm (left) and 5 µm (right). (**D**) Colocalization analysis of Nora virus signal with progenitor cells marked using the *esg-Gal4, Gal80^ts^* driver crossed to *UAS-GFP*. Intestines were fixed and stained for DNA (DAPI, blue) and Nora virus (Cy3, red) Scale bars, 2 µm (left) and 5 µm (right). (**E**) Representative confocal images of intestines from Nora-infected *GFP::Dicer-2* flies stained for DNA (DAPI, blue), actin (RFP, red), Dicer-2 (GFP, green), and Nora virus (Cy5, purple). Scale bars, 10 µm (left) and 5 µm (right). (**F**) Survival analysis of Nora-infected (+) and non-infected flies (-) following intestinal infection with *P. aeruginosa* PA14 (OD 0.25) or *S. marcescens* Db11 (OD 0.25), or exposure to 1 mM paraquat. (**G**) Nora virus RNA levels measured by RT-qPCR in flies from the experimental conditions shown in (F). (**H**) Representative confocal images of intestines from Nora-infected flies following PA14 infection, Db11 infection, or paraquat exposure. Tissues were fixed and stained for DNA (DAPI, blue), actin (FITC, green), and Nora virus (Cy3, red). Scale bar, 3 µm. (**I**) Quantification of (H). EC: enterocyte; ISC: progenitor cells. Each graphic is representative of three independent experiments. Survival panels are presented as mean ± SEM with each curve representing a triplicate of 20 flies. Other panels are presented as box and whiskers where the middle bar of the box plots represents the median, and the upper and lower limits of boxes indicate the first and third quartiles, respectively; the whiskers define the minima and maxima. Relative unit was used for Nora virus load panels. Each dot in Nora virus load panels (A, G) represent a sample of 5 flies. Survival data were analyzed using a log-rank (Mantel-Cox) test. Nora virus load was analyzed using a Mann-Whitney nonparametric test (A). Comparisons of Nora virus load under pathogen infection or paraquat exposure (G) and of quantification of Nora (+) ECs (I) were analyzed using one-way ANOVA with Tukey’s post-hoc test. Statistical significance is indicated as *p < 0.05, **p < 0.01, ***p < 0.001, ****p < 0.0001.

We have shown above that young Nora-positive flies are more susceptible to *P. aeruginosa* ingestion. We found that the Nora-positive flies were also more susceptible to the strongly oxidizing agent paraquat in survival experiment (Fig. 5F). As for *P. aeruginosa* oral infection, the ingestion of paraquat led to similarly enhanced levels of the viral load as monitored by RTqPCR (Fig. 5G) and by staining with the Nora antibody (Figs. 5H & S5D). Immunohistochemistry further revealed that the virus starts also proliferating within enterocytes under pathogenic or chemical stress (Fig. 5H-I). Interestingly, aged flies raised on our standard food or young flies raised on rich food also exhibited a growth of the virus in enterocytes (Fig. S5E-I), in keeping with the increased viral burden found in these 30-day-old flies (Fig. 3A-B). In addition, the detection of the Nora virus in enterocytes (Fig. 5I & Fig. S5I) also correlated with an enhanced proliferation of ISCs (Figs. 3C & S3D).

In conclusion, the Nora virus is initially restricted to progenitor cells in basal conditions but appears to start proliferating also inside enterocytes upon stress and/or increased proliferation of ISCs. Importantly, Nora-positive cells were detected only in the posterior midgut (scheme in Fig. S5J). These observations open the possibility that Nora virus is transmitted to differentiating cells after ISC division (Figs. 5C & S5B) and then ultimately to enterocytes.

### ISC division promotes the proliferation of the Nora virus

The above conclusion implies that ISC proliferation is likely needed to promote the proliferation of the virus. To test this hypothesis, we used two complementary approaches that rely on the fact that enterocyte damage triggers the compensatory proliferation of ISCs.

First, we measured the degree of apoptosis in epithelial cells through ApopTag/TUNEL staining. We noted that Nora-positive bacteria do not display enhanced levels of apoptosis in this tissue under basal conditions (Fig. 6A, Fig. S6A). However, the ingestion of *P. aeruginosa* led to enhanced levels of apoptosis, significantly more so in Nora-positive flies (Fig. 6A, Fig. S6B). In contrast, paraquat induced much higher levels of positive signals, independently of the Nora status of the exposed flies (Fig. 6A, Fig. S6C). This situation was roughly mirrored in the number of mitotic ISCs (Fig. 6B), although the degree of proliferation induced by paraquat was similar to that induced by *P. aeruginosa* infection, possibly mirroring an adverse effect of paraquat exposure to the division of ISCs (Chatterjee and Ip, 2009).

**Figure 6.**
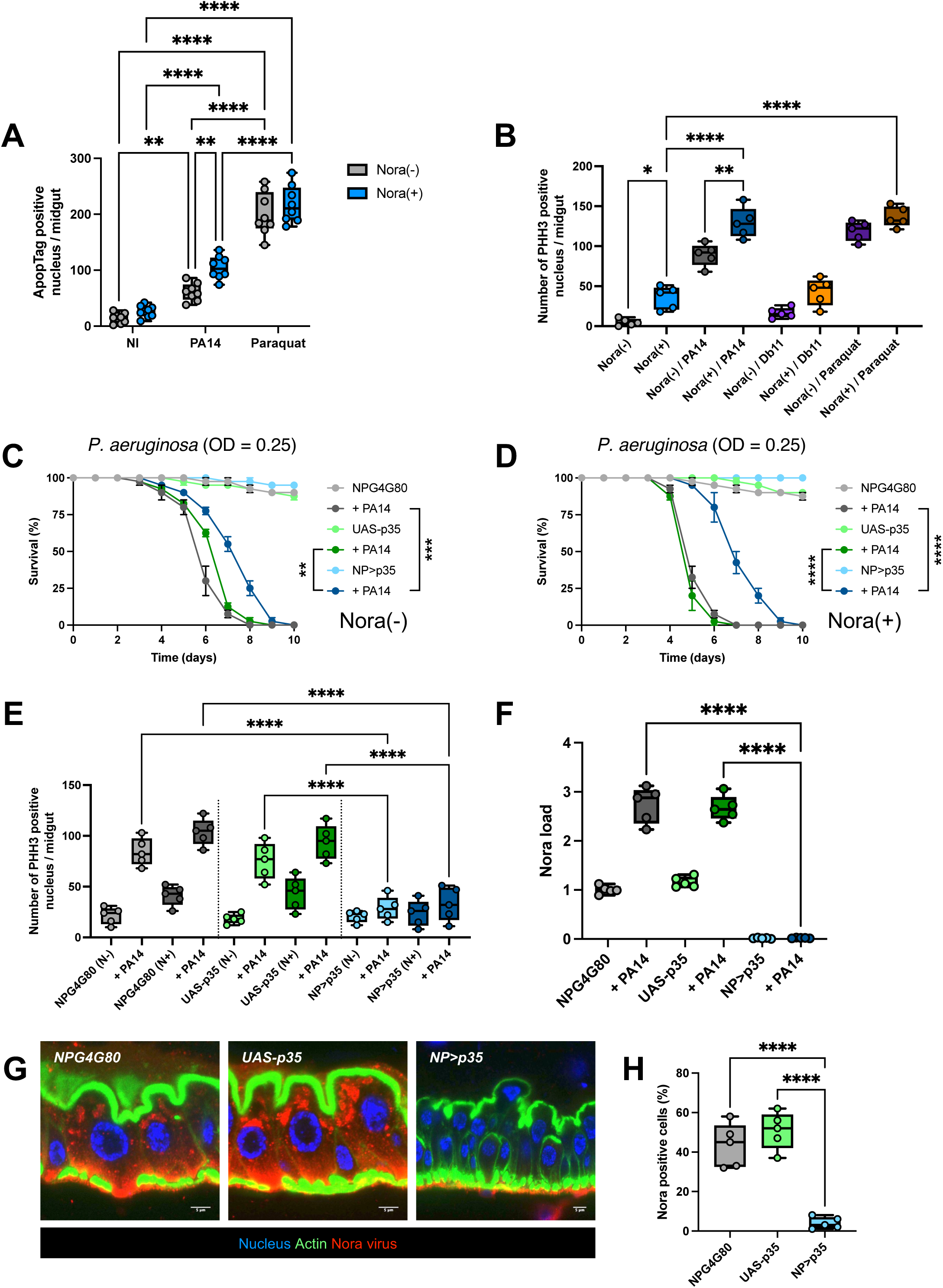
Inhibition of apoptosis prevents epithelial turnover and the proliferation of Nora virus. (**A**) Quantification of apoptotic nuclei in the posterior midgut (region R4 & R5 [see Fig. S5J]) using ApopTag staining following *P. aeruginosa* PA14 infection or 1 mM paraquat exposure and controls. (**B**) Quantification of phospho-histone H3 (PHH3)-positive nuclei in the posterior midgut of Nora-infected flies 24h after PA14 infection or paraquat exposure. (**C**) Survival analysis of non-infected flies (Nora(-)) following PA14 intestinal infection (+ PA14) in control genotypes (NPG4G80 driver: [*NP1-Gal4, Gal80^ts^*] and *UAS-p35* alone) and in flies with enterocyte expression of the apoptosis inhibitor gene *p35* (*NP>p35*). (**D**) Survival analysis of Nora-infected flies (Nora(+)) following PA14 intestinal infection in control genotypes and in flies expressing the *p35* gene in the intestine. (**E**) Quantification of PHH3-positive nuclei in the posterior midgut at 5 days following PA14 infection in flies from panels (C) and (D). PHH3-positive nuclei were quantified in control and apoptosis-inhibited intestines. (**F**) Nora virus RNA levels measured by RT-qPCR at 3 days following PA14 infection in intestines from flies shown in (D). (**G**) Representative confocal images of old intestines (30-day-old) from Nora-infected flies in control genotypes (*NP* driver (NPG4G80) and *UAS-p35* alone) and in flies expressing p35 in the intestine (*NP>p35*). Tissues were fixed and stained for DNA (DAPI, blue), actin (FITC, green), and Nora virus (Cy3, red). Scale bar, 5 µm. (**H**) Quantification of Nora virus-positive cells in the intestine from the experimental conditions shown in (G). Each graphic is representative of three independent experiments. Survival panel is presented as mean ± SEM with each curve representing a triplicate of 20 flies. Other panels are presented as box and whiskers where the middle bar of the box plots represents the median, and the upper and lower limits of boxes indicate the first and third quartiles, respectively; the whiskers define the minima and maxima. Relative unit was used for Nora virus load panels. Each dot in Nora virus load panel (F) represents a sample of 5 flies. Each dot in the PHH3 quantification panels (B, E) represent one single posterior midgut. Each dot in the ApopTag quantification panel (A) represents one single posterior midgut. Each dot in the Nora virus quantification panel (H) represents a region of one single posterior midgut. Survival data were analyzed using a log-rank (Mantel-Cox) test. ApopTag, PHH3, and Nora virus quantifications were analyzed using one-way ANOVA with Tukey’s post-hoc test (A, B, E, F, H). Statistical significance is indicated as *p < 0.05, **p < 0.01, ***p < 0.001, ****p < 0.0001.

Next, we attempted to block the induction of apoptosis in adult enterocytes by expressing there the baculovirus p35 protein that inhibits executioner caspases (Hay et al., 1994). p35 did partially protect the flies from the noxious effects of *P. aeruginosa* ingestion, with a relatively stronger effect on Nora-positive flies, which correlates with the higher number of apoptotic cells measured in those flies (Fig. 6C-D, Fig. 6A). As expected, the expression of p35 in enterocytes blocked the compensatory proliferation of ISCs (Fig. 6E). Strikingly, Nora virus was obliterated in both control or *P. aeruginosa*-infected, originally Nora-positive, flies, as monitored by RTqPCR (Fig. 6F) and by immunohistochemistry in 30-day old flies (Fig. 6G-H) for which less than ten Nora-positive progenitor cells per midgut were detected at best. The second approach was to interfere with the signals that drive ISC proliferation upon enterocyte damage. JAK-STAT pathway activation in ISCs promotes their division (Buchon et al., 2009b; Cronin et al., 2009; Jiang et al., 2009). Silencing in ISCs the genes that encode either the JAK-STAT pathway receptor Dome or the STAT92E transcription factor also provided protection against ingested *P. aeruginosa*, in both Nora-positive and Nora-negative cells (Fig. 7A-B). Indeed, the *upd3* gene encoding one Dome ligand and the gene encoding the JAK-STAT-regulated inhibitor SOCS36E were induced in dissected midguts (crop and Malpighian tubules not included) by *P. aeruginosa* ingestion, again more strongly so in Nora-positive flies (Fig. 7C). This result was further corroborated using a transgenic fly line that expresses an UPD3-GFP reporter protein (Fig. 7D). As expected, silencing *Dome* or *STAT* abolished the compensatory proliferation of ISCs (Fig. 7E). Again, as for the ectopic expression of p35 in enterocytes, blocking JAK-STAT pathway signaling in ISCs prevented any proliferation of the Nora virus in midguts, whether infected orally with *P. aeruginosa* or not (Fig. 7F). Microscopic analysis of immunostained posterior midguts confirmed that when the JAK-STAT signaling pathway was blocked specifically in ISCs using a *Dome-Gal4* driver line, Nora virus remained confined to progenitor cells and was not detected in enterocytes (Fig. 7G-H).

**Figure 7.**
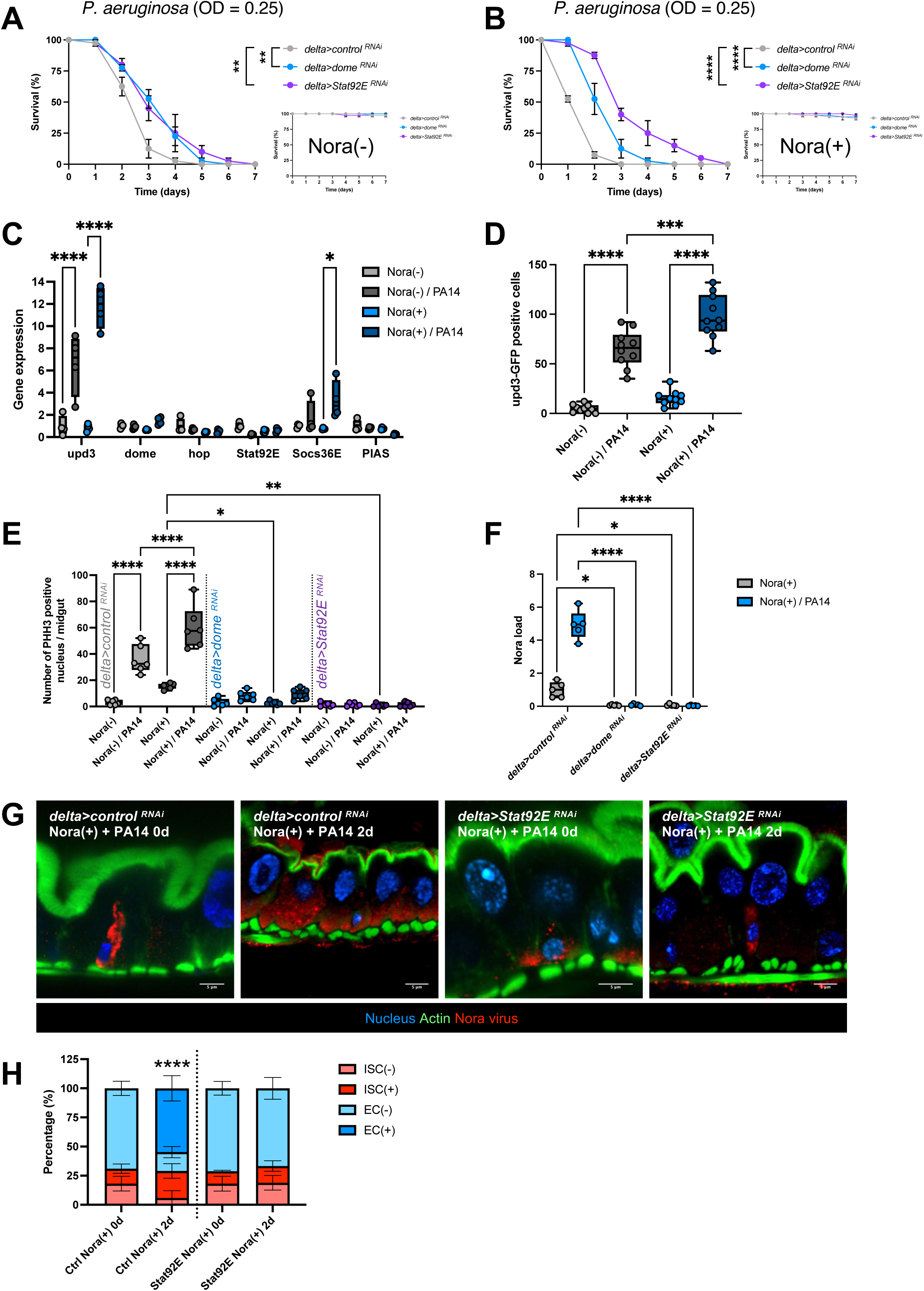
Inhibition of the JAK-STAT pathway prevents epithelial turnover and the proliferation of Nora virus. (**A-B**) Survival analysis of non-infected flies (A) and infected flies (B) following *P. aeruginosa* PA14 intestinal infection of flies in which RNAi-mediated silencing of JAK-STAT pathway component genes (*domeless* and *Stat92E*) in intestinal stem cells was achieved using the *Delta* driver. Control genotypes and non-infected (NI) survival assays are shown in the lower right insets. (**C**) Relative expression of JAK-STAT pathway-related genes (*upd3*, *dome*, *STAT92E*, *SOCS36E*, and *PIAS*) measured by RT-qPCR at 3 days following PA14 infection in Nora-infected (+) and non-infected (-) flies’ midguts. (**D**) Quantification of *upd3-GFP* reporter signal in the posterior midgut at 2 days following PA14 infection in Nora-infected (+) and non-infected (-) flies. (**E**) Quantification of phospho-histone H3 (PHH3)-positive nuclei in the posterior midgut at 3 days following PA14 infection in flies with RNAi-mediated silencing of *domeless* or *Stat92E* in intestinal stem cells using the *Delta-Gal4* driver. PHH3-positive nuclei were used as a measure of epithelial proliferation. (**F**) Nora virus RNA levels measured by RT-qPCR in intestines from flies shown in (E). (**G**) Representative confocal images of intestines from Nora-infected flies in control genotypes (*Delta>control RNAi*) and *UAS-p35* alone) and in flies expressing p35 in the intestine (*NP>p35*). Tissues were fixed and stained for DNA (DAPI, blue), actin (FITC, green), and Nora virus (Cy3, red). Scale bar, 5 µm. (**H**) Quantification of Nora-positive cells of the intestinal epithelium displayed in (G). EC: enterocyte; ISC: progenitor cells. Each graphic is representative of three independent experiments. Survival panels are presented as mean ± SEM with each curve representing a triplicate of 20 flies. Other panels are presented as box and whiskers where the middle bar of the box plots represents the median, and the upper and lower limits of boxes indicate the first and third quartiles, respectively; the whiskers define the minima and maxima. Relative unit was used for Nora virus load panels. Each dot in Nora virus load panel (F) represents a sample of 5 flies. Each dot in the PHH3 quantification panels (E) represent one single posterior midgut. Each dot in the gene expression panel (C) represents 5 posterior midguts. Each dot in the upd3-GFP quantification panel (D) represents one single posterior midgut. Survival data (A-B) were analyzed using a log-rank (Mantel-Cox) test. PHH3 quantification (E), gene expression analysis (C), upd3-GFP quantification (D), Nora virus load (F), and the number of Nora (+) enterocytes (H) were analyzed using one-way ANOVA with Tukey’s post-hoc test. Statistical significance is indicated as *p < 0.05, **p < 0.01, ***p < 0.001, ****p < 0.0001.

## Discussion

Nora virus infection is prevalent both in wild *D. melanogaster* flies and laboratory stocks (Habayeb et al., 2006; Webster et al., 2015). It has originally been reported to have limited effects on its host fitness, likely due to its proliferation that is mostly restricted to the intestinal epithelium (Habayeb et al., 2009b). Here, in keeping with some more recent work (Hanson and Lemaitre, 2023), we report that the presence of Nora virus may be a confounding factor when performing intestinal infections or exposing flies to xenobiotics, and more generally when investigating fitness in extended processes such as aging. Our data support a model according to which Nora virus primarily infects intestinal epithelium progenitor cells and largely remains latent/quiescent. However, exposure to a variety of stresses that all lead to an increased proliferation of ISCs, apparently reactivates the virus and results in the contamination of enterocytes. Ultimately, the generalized proliferation of Nora within epithelial cells affects the barrier function of the midgut epithelium and leads to a decreased fitness that leads to a premature demise of the infected flies (Fig. 8).

**Figure 8:**
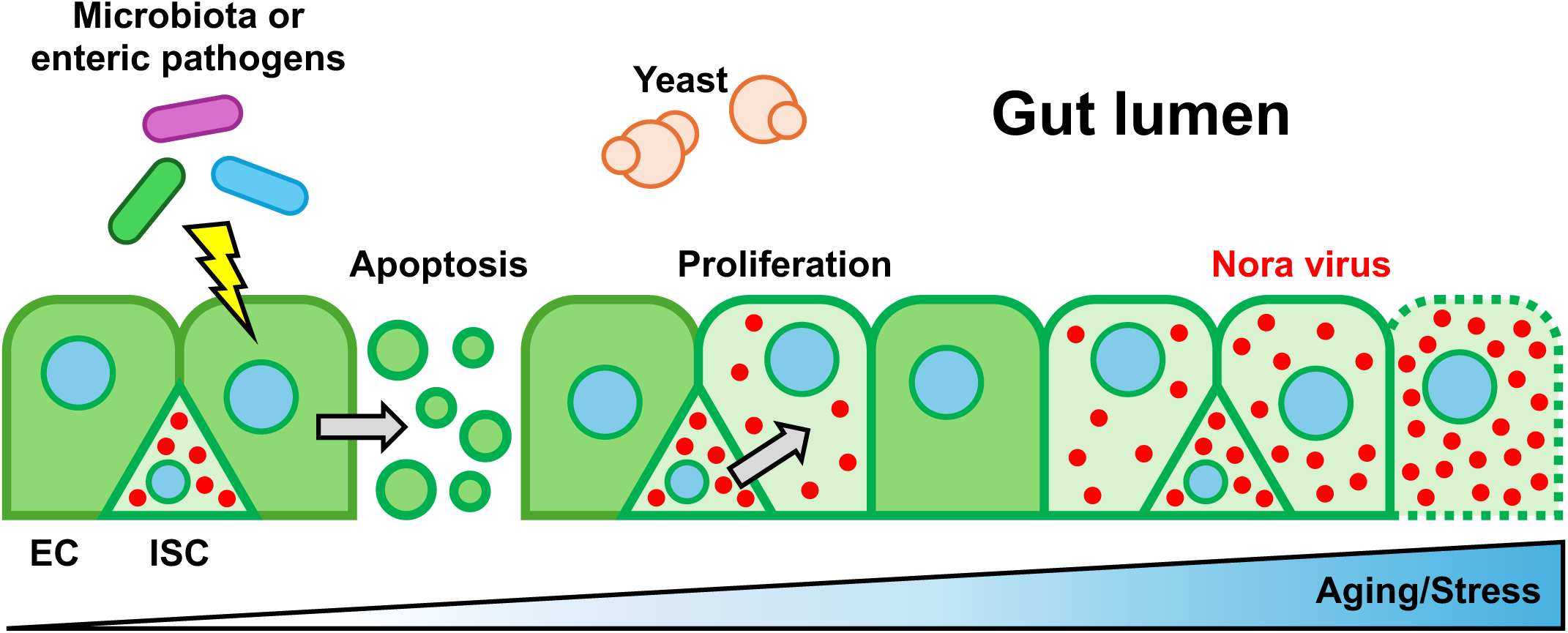
Model of Nora virus propagation in *Drosophila* intestine. Schematic representation of Nora virus propagation in the posterior midgut from intestinal stem cells to enterocytes. The Nora virus initially invades ISC in the midgut and remains rather quiescent. External factors such as age, infections or exposure to xenobiotics stress the host intestinal epithelium and induce ISC compensatory proliferation that ensures a degree of homeostasis by replacing damaged enterocytes. ISC cell division appears to stimulate the proliferation of the Nora virus that ultimately contaminates enterocytes. This is likely to further damage the epithelium and lead to a loss of the integrity of this intestinal barrier, allowing the enhanced passage of gut bacteria to the hemocoel.

Our data document a low basal level of infection in young flies, which correlates with a localization restricted to progenitor cells of the intestinal epithelium. Strikingly, a series of parameters such as age, food nutritional value, pathogenic infection of the gut, and a strong oxidizing agent all induce both an increased viral burden associated to the infection of enterocytes, and proliferation of ISCs. Nora virus appears either to have relatively moderate (Habayeb et al., 2009a; Hanson and Lemaitre, 2023; Nigg et al., 2024; Rogers et al., 2020; Schissel et al., 2021) or intermediate to strong (this study; (Hanson and Lemaitre, 2023)) effects on host fitness. The finding that the higher nutrient quality of the food leads to reduced effects on host fitness may partially account for these differences (Fig. 3D-E), even though the Nora load was much higher on nutrient-rich food (Fig. 3A). The genetic background may also contribute to these differences, even though we observed similar effects of Nora infection on host fitness in a variety of strains fed on sucrose solution (Fig. S1 E-I). It could also be that the strains used in other laboratories may have modifier genes in their genetic background; indeed, one of Oregon strains we tested appeared to be nonpermissive for Nora (Fig. S1I; see also (Hanson and Lemaitre, 2023)). Thus, the Nora load may differ in the different laboratories and could account for the differences in host fitness(Hanson and Lemaitre, 2023). Interestingly, the microbiota had opposite effects on host fitness depending on the food source (positive on sucrose (Fig. S3G) *vs.* negative on standard or rich food (Fig. 3D-E)) (see also Arias-Rojas and Iatsenko, 2022 and references therein). We did observe a higher microbiota titer in Nora-infected flies (Figs. 3C, S3F), which however did not correlate with increased proliferation of ISCs at least on sucrose (Fig. S3H). Of note, strains kept in our laboratory do not contain *Acetobacter* but mostly *Lactobacilli* strains in their microbiota. Whether there is an increased diversity of the microbiota remains to be addressed.

Our results thus suggest that the virus is activated in dividing progenitor cells, especially when the intestinal epithelium is stressed either by infection or exposure to xenobiotics, and goes on proliferating in differentiated cells derived from these ISCs. The rate of ISC proliferation is always higher in Nora-positive flies, suggesting that Nora contamination of enterocytes may contribute in addition to stressors to their apoptosis and consequently lead to an enhanced compensatory proliferation of ISCs. The finding that blocking apoptosis enhances the fitness of Nora-positive flies to *P. aeruginosa* infection to the level of Nora-negative flies while equally decreasing the proliferation of ISCs supports this proposition (Fig. 6C-E). Hence, whenever ISC proliferation is triggered, the additional death of enterocytes caused by Nora virus amplifies the phenomenon in a positive feedforward loop. This model is further strengthened by the finding that the JAK-STAT ligand UPD3 is expressed at a higher basal level and induced more strongly by *P. aeruginosa* infection in Nora-positive flies than in Nora-negative flies. Thus, blocking either enterocyte apoptosis or the regulatory pathway that drives compensatory ISC proliferation yields a similar protection against the effects of Nora infection on fly fitness. More importantly, both experimental manipulations lead to a spectacular decrease of the viral burden, even under *P. aeruginosa* infection conditions (Fig. 6F & Fig. 7F).

We initially detect Nora-positive cells in basally located diploid progenitor cells. This observation begs two questions. The first one is how basally located cells can become infected and the second is why enterocytes that make up the apical part of the epithelium do not get initially infected. The second question will be discussed further below. With respect to the first one, we cannot formally exclude that enterocytes transport viral particles to the basal side where they would then infect the progenitor cells. Indeed, many arboviruses cross the intestinal barriers and transit through enterocytes in a polar manner (Hodgson et al., 2024). Alternatively, progenitor cells are known to extend thin processes, some of which might reach and probe luminal contents (Ohlstein and Spradling, 2006). These processes might carry unidentified receptors for the Nora virus and enable the primary infection of progenitor cells. In this view, it is an open possibility that enterocytes do not express these putative receptors.

Our experiments point to an important role of initial replication of Nora virus in ISCs. With the exception of hematopoietic stem cells that can be infected by retroviruses and herpes viruses, mammalian germ cells, embryonic and adult stem cells have long been known to be rather resistant to viral infections (Wu et al., 2019). Interestingly, mammalian stem cells do not appear to rely primarily on interferon-based defenses, even though they do express a subset of interferon-stimulated genes (ISGs) that contribute to the defense against viral infections (Wu et al., 2018). Yet, they are not responsive to interferon, which is mediated through JAK-STAT signaling, and instead rely to a large extent on RNA interference (Maillard et al., 2013; Maillard et al., 2016; Poirier et al., 2021). Whereas interferon signaling in mammals may have adverse effects on stem cell function because of antiproliferative actions of some ISGs, JAK-STAT signaling in *Drosophila* ISCs promotes stem cell division and is also involved in the subsequent differentiation of progenitor cells (Jiang et al., 2009). Interestingly, picornaviruses and coxsackieviruses are known to productively infect activated or proliferating cells while the infection of quiescent cells leads only to persistence or latency (Feuer and Whitton, 2008). Future studies will tell at which step of the cell cycle the Nora virus preferentially proliferates.

We have not addressed here whether the Nora virus is susceptible to host intestinal antiviral defenses. A major antiviral defense in *Drosophila* is the RNAi pathway (Galiana-Arnoux et al., 2006; van Rij et al., 2006), which however appears less active in the gut (Mondotte et al., 2018), albeit oral DCV infection led to an increased viral load in *Ago2* but not *Dcr2* mutant (Segrist et al., 2021). It will be interesting to determine whether the absence of a Dcr2-fluorescent proteins fusions in progenitor cells that we report in this study rules out a role for the RNAi pathway in intestinal host defense against the Nora virus. Nora virus-derived siRNAs have been detected in fly tagmata, including the head (van Mierlo et al., 2012). In addition, Nora VP1 protein has been shown to inhibit slicer activity of Ago2 (van Mierlo et al., 2012). In contrast, it has been reported that Nora virus load is not altered in flies defective for the RNAi antiviral pathway (Habayeb et al., 2009b), although the same study failed to see an effect of the JAK-STAT pathway, which conflicts with our own results (this study). It should be emphasized that this study was performed on young 3-5day old flies in which Nora virus is detected only in progenitor cells and not in enterocytes. Thus, it is an open possibility that the RNAi machinery is efficient in preventing the primary infection of enterocytes (but would not impair polarized transit to the basal side of enterocytes). Consequently, only enterocytes derived from infected progenitor cells would allow the contamination of enterocytes. Possibly, a high load of the virus in progenitor cells would allow it to interfere with the RNAi machinery in developing enterocytes. The PVR/ERK pathway has been documented to play a role in enterocyte host defense against viral infections (Sansone et al., 2015). However, it needs to be primed by the microbiota through the PGRP-LC/IMD pathway and Pvf2 induction in enterocytes. Given the opposite effects of the microbiota on Nora-infected fly fitness depending on the food source, it appears unlikely that the PVR/ERK signaling pathway plays an essential role in host defense against the Nora virus. In addition, the likely absence of *Acetobacter* species in the microbiota of our fly strains on our food likely limits the induction of Pvf2 expression in enterocytes via the IMD pathway since *Lactobacilli* do not induce it (Sansone et al., 2015). The STING/Relish pathway is another important systemic antiviral pathway (Ai et al., 2024; Cai et al., 2020; Goto et al., 2020; Segrist et al., 2024), which is also relevant to that of the intestinal epithelium. However, current data suggest it is mostly active in enterocytes (Nigg et al., 2024; Segrist et al., 2021). Importantly, in the case of DAV, another prevalent intestinal virus, it is the EGFR pathway in enterocytes and not the JAK-STAT pathway acting in progenitor cells that is important for DAV proliferation. In addition, DAV is mostly found in enterocytes and only occasionally in progenitor cells (Nigg et al., 2024). Thus, while this axis of defense may not be essential once stem cells are proliferating, it might protect the enterocytes in the primary fecal infection.

As emphasized by other investigators, intestinal viruses may be confounding factors of studies on aging and intestinal physiology, especially as regards the proliferation of ISCs (Hanson and Lemaitre, 2023; Nigg et al., 2024). Our results further add intestinal bacterial infections and exposure to toxicants to this list and more generally any study in which the process under investigation involves monitoring the survival of flies for periods of time over eight days. Finally, we have attempted to establish a Nora-free facility and bleached some 100 fly lines that tested Nora-negative on whole fly extracts. However, after a few months many of these lines tested positive for Nora. It is an open possibility that some fly strains cannot be cured (we had to go through several rounds of bleaching to cure from Nora contamination our Ore-R (SM) stock). Alternatively, a more sensitive test for Nora detection should be used, starting on dissected guts and for instance digital PCR (see also (Nigg et al., 2022)).

## Materials & Methods

### Drosophila stock and husbandry

Wild-type Ore-R(SM) Nora(+) and Ore-R(SC) Nora(-) fly stocks were found in our laboratory. Both stocks of wild-type *Oregon-R* flies tested negative for *Wolbachia* infection. Stock used for the septic injury and natural infections are the following: wild-type *w^A5001^* (Thibault et al., 2004), mutant *MyD88c03881* (Tauszig-Delamasure et al., 2002), mutant DD1 *cn bw* (Rutschmann et al., 2000), *key^cO2831^*(Ferrandon D., unpublished), *NP1-Gal4* (Cronin et al., 2009), *esg-Gal4Gal80^ts^*(Micchelli and Perrimon, 2006), *delta-Gal4* (Zeng et al., 2010), VDRC GD control line (#6000 VDRC), *UAS-dome^RNAi^* (#36355 GD VDRC) (Borensztejn et al., 2013), Bloomington control line *UAS-mCherry^RNAi^* (#35787 Bloomington Drosophila Stock Center) (Borensztejn et al., 2013), *UAS-p35* (#5073 Bloomington Drosophila Stock Center) (Hay et al., 1994; Rahmatika et al., 2019), *UAS-Stat92E^RNAi^*(#26899 Bloomington Drosophila Stock Center) (Nagai et al., 2023), and *upd3::GFP* (Zhou et al., 2013).

All stocks have been checked for the presence of known pathogens and symbiont besides Nora virus (Haller et al., 2014; Lestradet et al., 2014; Niehus et al., 2012). Fly stocks were kept at 25°C and nearly 60% humidity, on a standard semi-solid cormeal medium (6.4% (w/v) cornmeal (Moulin des moines, France), 4.8% (w/v) granulated sugar (Erstein, France), 1.2% (w/v) yeast brewer’s dry powder (VWR, Belgium), 0.5% (w/v) agar (Sobigel, France), 0.004% (w/v) 4-hydroxybenzoate sodium salt (Merck, Germany)). For lifespan analysis, 3 x 20 female flies were kept at 25°C with 60% humidity, on standard fly food. Flies were transferred without anesthesia on fresh food every 4 days. For survival test on a sucrose only diet, 3x 20 female flies were fed on 2 mL on a 50 mM sucrose solution and kept at 25°C with 60% humidity.

### Microbial strains, growth conditions and infection

Microbes were grown in these conditions: *Pseudomonas aeruginosa* PA14 in Brain-Heart-Infusion Broth (BHB), overnight, at 37°C (Rahme et al., 1995); *Serratia marcescens* Db11 (Nehme et al., 2007) in Luria Bertoni Broth (LB), overnight, at 37°C; *Enterobacter cloacae* (Lemaitre et al., 1997), in LB, overnight, at 30°C; *Enterococcus faecalis* (TX0016) in BHB overnight, at 37°C; *Beauvaria bassiana* (Lemaitre et al., 1997) on malt agar plates at 25°C. Intestinal infections were performed as described previously with PA14 (Haller et al., 2014) and Db11 (Lee et al., 2016). Briefly, flies were continuously feeding on a sucrose solution containing the bacteria at the indicated OD and deposited on filter pads in the absence of standard food. In the case of *S. marcescens* 100 µL of 100 mM sucrose solution was added to the cage every day. Infected and control flies were kept at 25°C. For PA14 infection, 3x 20 female flies were used per experiment. For *E. cloacae* septic injury, 50 mL of an overnight culture of bacteria was centrifuged 30 min at 3 000 x g. The supernatant was removed and pellet used to infect the flies. For *E. faecalis* septic injury, overnight culture was diluted to 1/50 in fresh BHB and allowed to growth for 3 more hours at 37°C. The final culture was centrifuged 10 min at 3000 x g and the pellet washed one time with sterile PBS. The bacteria were resuspended to a final OD = 0.5 that was used to infect female flies with a septic injury. 3x 20 females were infected with each bacterium. The septic injury and the PBS clean injury were performed as described (Haller et al., 2014): an electrolysis-sharpened tungsten wire dipped or not in a microbial solution was used to prick flies in the thorax at a position corresponding to alary muscles. 20 females were injured. For *B. bassiana* natural infection, flies were anesthetized, deposited and shaken on a sporulating plate containing the fungus. 3 x 20 females were infected. Flies were kept at 29°C with 60% humidity and transferred, without anesthesia, to fresh food every 2-3 days.

### Paraquat exposure

Paraquat solution was prepared by dissolving Paraquat dichloride (Sigma-Aldrich # 50636) in 50 mM sucrose solution to obtain a final paraquat concentration at 1 mM. For survival test on a paraquat supplemented sucrose only diet, 3x 20 female flies were fed on 2 mL on a 50 mM sucrose plus 1 mM paraquat solution and kept at 25°C with 60% humidity.

### Dechorionation of *Drosophila* eggs

Nora(+) flies were allowed to lay eggs overnight at 25°C in a cage on apple juice agar plate with yeast paste in the center of the plate. Eggs were collected, washed with water, and dechorionated with a 50% bleach solution for 3 min with constant up and down pipetting of the solution. Eggs were abundantly rinsed with water, aligned under the microscope on a piece of agar medium and transferred by capillarity on a coverslip. One drop of mineral oil was applied to cover the eggs, and the coverslip was deposited on a petri dish with normal *Drosophila* food. After 2 days, larvae were transferred to normal food vial (single bleaching) or treated again with bleach solution as previously and then transferred to normal food vial (double bleaching or even triple bleaching if needed). Once flies emerged, they were tested for Nora virus infection. Usually, there was no difference in the results obtained with the single or double bleaching procedures although this observation applies only to short term observations.

### Pure Nora virus suspension preparation

Nora(+) flies (around 5 000 flies) were crashed in ice cold PBS using a potter homogenizer. The homogenate (around 35 mL) was transferred in a 50 mL Falcon tube and centrifuged at 1 500 rpm for 10 min, at 4°C in an Eppendorf 5810R centrifuge to remove wings, legs and other fly debris. The resulting supernatant was sequentially filtrated through 0.8, 0.45 and 0.22 μm filter units and stored at −80°C. The homogenate (around 26 mL) was then layered in two 15 ml Beckmann Ultra-Clear centrifuge tubes over 1.5 mL of a 30 % (wt/wt) sucrose solution (50 mM Hepes pH 7.0, 0.1 % BSA) and centrifuged at 24,000 rpm for 6.5 hours at 11°C using a JS-24.15 rotor (Beckmann). The pellets were resuspended in a total volume of 500 μL Hepes solution (Hepes 50 mM, pH 7.0) and transferred into a 1.5 mL Eppendorf tube. The solution was then centrifuged at 10,000 rpm for 10 min in a 5424 Eppendorf centrifuge to discard insoluble material. The resulting supernatant containing virus particles was layered over a 40-10 % (wt/wt) sucrose gradient (50 mM Hepes, pH 7.0) prepared in a 15 mL Ultra-Clear centrifuge tube (Beckmann) and centrifuged at 11°C for 4h at 24,000 rpm in a JS-24.15 rotor (Beckmann). An opalescent band, that migrated near the middle of the tube, containing virus particles was then collected by puncturing the tube with a 25-gauge needle mounted on a 1 ml syringe. This solution was then layered onto a 30 % (wt/wt) sucrose solution contained in a 15 mL Beckmann Ultra-Clear centrifuge tubes (50 mM Hepes pH 7.0, 0.1 % BSA) and centrifuged at 24,000 rpm for 6.5 hours at 11°C using a JS-24.15 rotor (Beckmann). The pellet was resuspended in 500 μl of Tris solution (Tris-HCl 10 mM, pH 7.5), aliquoted, and stored at −80°C.

### Nora virus electronic microscopic picture

The sample was prepared by plunge-freezing of 2.5 μL of the sample on a holey carbon Quantifoil R 2/2 grid using FEI Vitrobot Mark IV machine. Images were gained on FEI Polara F30 TEM microscope with acceleration voltage 100 KV and underfocus around 2.0 microns.

### Nora virus re-infection with the pure viral preparation

Nora(-) flies were fed 24 hours with a 1/100 dilution of the viral preparation in sucrose 50mM. In practice, 200 μL of the 1/100 dilution (in sucrose 50mM) of the viral suspension were deposited in an Eppendorf cup that was then placed in an empty fly tube. Flies (male and females) were then added in the tube. The flies were allowed to feed on the viral suspension for 24 hours and then transferred to a fly tube containing standard fly food.

### Nora virus re-infection with feces

200 males of Nora(+) flies were allowed to deposit their feces in a food vial for 5 days at 25°C. Nora(+) flies were then replaced by 50 males and 50 females of cured Nora(+) flies. After 5 days, the vial was emptied, and Nora virus infection status was monitored in those flies (generation G0). Once the progeny emerged, 0 to 4 days old flies were transferred to a fresh vial for 4 days and then monitored for Nora virus load or used for experiments (first generation after re-infection G1). The same procedure was repeated with the second generation after re-infection (G2).

### Generation of a polyclonal antibody that detects the Nora virus

Pure virus preparation was mixed with complete (first inoculation) then incomplete Freund’s adjuvant and was injected intraperitoneally in BALB/c mice every week over a period of three weeks prior to ultimately bleeding the mice. The specificity of the antibody was assessed by immunohistochemistry on Nora-negative and Nora-positive flies as determined using RTqPCR.

### Immunostainings

Midguts were dissected in PBS and fixed for 30 minutes with 4% paraformaldehyde. Samples were washed three times with PBS-Triton X-100 0.1% (PBT 0.1%) as described (Socha et al., 2023). For actin staining midguts were incubated for 1h30 at room-temperature or overnight at 4°C in 10 μM Fluorescein Isothiocyanate (FITC) (Sigma-Aldrich #P5282) or Texas-Red labeled phalloidin (Invitrogen TM #T7471). Samples were then washed three times with PBT 0.1%. All samples were mounted on diagnostic microscope slides (Thermo Fisher Scientific) with Vectashield plus DAPI (Vector Laboratories). Samples were observed using a LSM780 confocal microscope (Zeiss) or in Axioskop 2 microscope (Zeiss). All images were analyzed with the ImageJ/Fiji software.

### Phospho-histone H3 immunostaining and microscopy

Fly guts were dissected, fixed, stained by immunohistochemistry with an anti-pH3 antibody (Millipore) and mounted following standard procedures (Lee et al., 2016). Midguts were observed using a fluorescent microscope (Axioskop 2, ZEISS) and nuclei positive for pH3 staining were counted manually.

### Microbiota quantification

Female flies were sterilized for 30 seconds in 70% ethanol and midguts were dissected in sterile PBS (10 midguts per sample) and immediately transferred in sterile PBS. Midguts were homogenized with a sterile pestle. Serial dilutions of the suspension were performed prior to plating on specific solid media and incubated at 30°C. *Acetobacteriaceae* plates, *Enterobacteriaceae* plates and MRS plates were performed as described (Guo et al., 2014). For BHB agar plates: 37 g/L BHB (Sigma), 15 g/L agar (Sigma). Media were autoclaved 15 min, at 121°C (prior to use) and stored at 4°C.

### Bacterial titer in the hemolymph

PA14 presence in the hemolymph was assessed as described previously (Haller et al., 2014); briefly, hemolymph was collected by capillarity using a drawn-out pipette mounted on the Nanoject II microinjector in the absence of oil. Hemolymph from 10 female flies per sample was used.

### RT-qPCR analysis of gene expression in *Drosophila* midgut

Fly midguts (without crop, hindgut and Malpighian tubules) were dissected (20 per sample) and RNAs were extracted in TRI Reagent®-RT as described (Lee et al., 2016). Reverse transcription was performed using iScriptTM (BIO-RAD). Quantitative PCR was performed using iQTM SYBR® Green (BIO-RAD) and a C1000TM Thermal Cycler (BIO-RAD) device. Expression values were calculated using a standard curve (with genomic DNA) and normalized to *rp49* expression level. Results presented the average ± SD of 3 independent experiments (with biological duplicates or triplicates). See primer sequences in Table 1.

### RT-qPCR analysis of Nora virus load

Whole flies (6 per sample) were frozen at −80°C. RNA extraction with a Nucleospin® RNA kit (Macherey-Nagel), reverse transcription and quantitative PCR were performed as described (Haller et al., 2014). Quantification values were calculated using a standard curve obtained from serial dilutions of a plasmid carrying the Nora virus DNA sequence amplified by the couple of primers used for the PCR. See primer sequences in Table 1.

To check the absence of other potentially contaminating viruses in the purified Nora virus preparations, we used the set of primers described by (Wu et al., 2010).

### Antibiotics treatment

3x 20 female flies (3-8 days old) were kept on a sucrose (50mM) only diet with addition of a 5 antibiotics cocktail: 100 μg/mL ampicillin, 50 μg/mL vancomycin, 100 μg/mL neomycin, 100 μg/mL metronidazole and 50 μg/mL tetracyclin. Tubes were changed every 3 days and flies transferred without anesthesia.

### SMURF

The SMURF assay was developed to observe the integrity of the intestinal barrier (Rera et al., 2011). Flies are kept on a sucrose solution colored with Blue food dye (Brilliant Blue FCF E133). After ingestion, if there is an intestinal integrity loss, the dye will diffuse in the hemolymph, coloring the fly in blue.

### ApopTAG

Used as described by the manufacturer Sigma-Aldrich for the product ApopTag Fluorescein In Situ Apoptosis Detection Kit.

### JAK-STAT gene expression

Fly midguts (without crop, hindgut and Malpighian tubules) were dissected (20 per sample) and RNAs were extracted in TRI Reagent®-RT as described (Lee et al., 2016). Reverse transcription was performed using iScriptTM (BIO-RAD). Quantitative PCR was performed using iQTM SYBR® Green (BIO-RAD) and a C1000TM Thermal Cycler (BIO-RAD) device. Expression values were calculated using a standard curve (with genomic DNA) and normalized to *rp49* expression level. Results presented the average ± SD of 3 independent experiments (with biological duplicates or triplicates).

### Statistical analysis and reproducibility

All statistical analyses were performed on Graphpad Prism version 10 (Graphpad software Inc., San Diego, CA). The Mann–Whitney tests were used unless otherwise indicated. For survival experiments, the time it takes for 50% of the flies to succumb (LT50) was determined for each curve being compared. An unpaired *t*-test used on the LT50s from biological triplicates was used to assess the significance between two survival curves. When using parametric tests (analysis of variance (ANOVA) and *t*-test), a Gaussian distribution of data was checked using either D’Agostino-Pearson omnibus, Anderson-Darling, or Shapiro-Wilk normality tests. All experiments were performed at least three times. Significance values: \**p* < 0.05, \*\**p* < 0.01, \*\*\**p* < 0.001, \*\*\*\**p* < 0.0001.

## Acknowledgements

We thank Pr. JL. Imler for continuous support of this project, Dr. S. Limmer, Dr. A. Ayyaz, and J. Bourdeaux for early help with experiments, MC. Lacombe, and E. Santiago for expert technical assistance. We are also grateful to Dr. D. Rodrigues for checking the stability of ectopically expressed GFP in ISCs. SH has been funded by the University of Strasbourg (Présidence fellowship) and by a FRM 4th year Ph. D fellowship (FDT20130928220). VB was supported by a fellowship from MENRT. Dominique Ferrandon was leading a Équipe Fondation Recherche Médicale (FRM DEQ20090515394). This work has also been funded by Institutional funds from CNRS and by ANR (DROSOGUT), NIH (PO1 AI070167), the Investissement d’Avenir program Laboratoire d’Excellence (NetRNA ANR-10-LABX-36; I2MC ANR-11-EQPX-0022).

## Notes

### Competing Interest Statement

The authors have declared no competing interest.

### Summary of Updates

Corrections made based on eLife reviewers comments.

## References

Ai, X., Deng, H., Li, X., Wei, Z., Chen, Y., Yin, T., Zhang, J., Huang, J., Li, H., Lin, X., et al. (2024). cGAS-like receptors drive a systemic STING-dependent host response in Drosophila. Cell reports 43, 115081.

Apidianakis, Y., Pitsouli, C., Perrimon, N., and Rahme, L. (2009). Synergy between bacterial infection and genetic predisposition in intestinal dysplasia. Proc Natl Acad Sci U S A 106, 20883–20888.

Arias-Rojas, A., and Iatsenko, I. (2022). The Role of Microbiota in Drosophila melanogaster Aging. Front Aging 3, 909509.

Ayyaz, A., and Jasper, H. (2013). Intestinal inflammation and stem cell homeostasis in aging Drosophila melanogaster. Front Cell Infect Microbiol 3, 98.

Biteau, B., Hochmuth, C.E., and Jasper, H. (2008). JNK activity in somatic stem cells causes loss of tissue homeostasis in the aging Drosophila gut. Cell Stem Cell 3, 442–455.

Bonfini, A., Liu, X., and Buchon, N. (2016). From pathogens to microbiota: How Drosophila intestinal stem cells react to gut microbes. Dev Comp Immunol 64, 22–38.

Borensztejn, A., Boissoneau, E., Fernandez, G., Agnes, F., and Pret, A.M. (2013). JAK/STAT autocontrol of ligand-producing cell number through apoptosis. Development 140, 195–204.

Broderick, N.A., and Lemaitre, B. (2012). Gut-associated microbes of Drosophila melanogaster. Gut Microbes 3, 307–321.

Buchon, N., Broderick, N.A., Chakrabarti, S., and Lemaitre, B. (2009a). Invasive and indigenous microbiota impact intestinal stem cell activity through multiple pathways in Drosophila. Genes Dev 23, 2333–2344.

Buchon, N., Broderick, N.A., Poidevin, M., Pradervand, S., and Lemaitre, B. (2009b). Drosophila intestinal response to bacterial infection: activation of host defense and stem cell proliferation. Cell Host Microbe 5, 200–211.

Buchon, N., Osman, D., David, F.P., Fang, H.Y., Boquete, J.P., Deplancke, B., and Lemaitre, B. (2013). Morphological and molecular characterization of adult midgut compartmentalization in Drosophila. Cell Rep 3, 1725–1738.

Cai, H., Holleufer, A., Simonsen, B., Schneider, J., Lemoine, A., Gad, H.H., Huang, J., Huang, J., Chen, D., Peng, T., et al. (2020). 2’3’-cGAMP triggers a STING- and NF-kappaB-dependent broad antiviral response in Drosophila. Sci Signal 13.

Chakrabarti, S., and Visweswariah, S.S. (2020). Intramacrophage ROS Primes the Innate Immune System via JAK/STAT and Toll Activation. Cell Rep 33, 108368.

Chatterjee, M., and Ip, Y.T. (2009). Pathogenic stimulation of intestinal stem cell response in Drosophila. J Cell Physiol 220, 664–671.

Chen, J., Lin, G., Ma, K., Guo, Y., Li, Z., Wang, X., and Ferrandon, D. (2025). N-acetyl-glucosamine primes Pseudomonas aeruginosa for virulence through a type IV pili/cAMP-mediated morphology transition. Nature communications 16, 9405.

Chen, J., Lin, G., Ma, K., Li, Z., Liegeois, S., and Ferrandon, D. (2024). A specific innate immune response silences the virulence of Pseudomonas aeruginosa in a latent infection model in the Drosophila melanogaster host. PLoS Pathog 20, e1012252.

Choi, N.H., Kim, J.G., Yang, D.J., Kim, Y.S., and Yoo, M.A. (2008). Age-related changes in Drosophila midgut are associated with PVF2, a PDGF/VEGF-like growth factor. Aging Cell 7, 318–334.

Cronin, S.J., Nehme, N.T., Limmer, S., Liegeois, S., Pospisilik, J.A., Schramek, D., Leibbrandt, A., Simoes Rde, M., Gruber, S., Puc, U., et al. (2009). Genome-wide RNAi screen identifies genes involved in intestinal pathogenic bacterial infection. Science 325, 340–343.

Donelick, H.M., Talide, L., Bellet, M., Aruscavage, P.J., Lauret, E., Aguiar, E., Marques, J.T., Meignin, C., and Bass, B.L. (2020). In vitro studies provide insight into effects of Dicer-2 helicase mutations in Drosophila melanogaster. RNA 26, 1847–1861.

Ekstrom, J.O., Habayeb, M.S., Srivastava, V., Kieselbach, T., Wingsle, G., and Hultmark, D. (2011). Drosophila Nora virus capsid proteins differ from those of other picorna-like viruses. Virus Res 160, 51–58.

Ekstrom, J.O., and Hultmark, D. (2016). A Novel Strategy for Live Detection of Viral Infection in Drosophila melanogaster. Sci Rep 6, 26250.

Ferrandon, D. (2013). The complementary facets of epithelial host defenses in the genetic model organism Drosophila melanogaster: from resistance to resilience. Curr Opin Immunol 25, 59–70.

Ferreira, A.G., Naylor, H., Esteves, S.S., Pais, I.S., Martins, N.E., and Teixeira, L. (2014). The Toll-dorsal pathway is required for resistance to viral oral infection in Drosophila. PLoS Pathog 10, e1004507.

Feuer, R., and Whitton, J.L. (2008). Preferential coxsackievirus replication in proliferating/activated cells: implications for virus tropism, persistence, and pathogenesis. Curr Top Microbiol Immunol 323, 149–173.

Galiana-Arnoux, D., Dostert, C., Schneemann, A., Hoffmann, J.A., and Imler, J.L. (2006). Essential function in vivo for Dicer-2 in host defense against RNA viruses in drosophila. Nat Immunol 7, 590–597.

Girardi, E., Lefevre, M., Chane-Woon-Ming, B., Paro, S., Claydon, B., Imler, J.L., Meignin, C., and Pfeffer, S. (2015). Cross-species comparative analysis of Dicer proteins during Sindbis virus infection. Sci Rep 5, 10693.

Goto, A., Okado, K., Martins, N., Cai, H., Barbier, V., Lamiable, O., Troxler, L., Santiago, E., Kuhn, L., Paik, D., et al. (2020). The Kinase IKKbeta Regulates a STING-and NF-kappaB-Dependent Antiviral Response Pathway in Drosophila. Immunity 52, 200.

Guo, L., Karpac, J., Tran, S.L., and Jasper, H. (2014). PGRP-SC2 promotes gut immune homeostasis to limit commensal dysbiosis and extend lifespan. Cell 156, 109–122.

Habayeb, M.S., Cantera, R., Casanova, G., Ekstrom, J.O., Albright, S., and Hultmark, D. (2009a). The Drosophila Nora virus is an enteric virus, transmitted via feces. J Invertebr Pathol 101, 29–33.

Habayeb, M.S., Ekengren, S.K., and Hultmark, D. (2006). Nora virus, a persistent virus in Drosophila, defines a new picorna-like virus family. J Gen Virol 87, 3045–3051.

Habayeb, M.S., Ekstrom, J.O., and Hultmark, D. (2009b). Nora virus persistent infections are not affected by the RNAi machinery. PLoS One 4, e5731.

Haller, S., Franchet, A., Hakkim, A., Chen, J., Drenkard, E., Yu, S., Schirmeier, S., Li, Z., Martins, N., Ausubel, F.M., et al. (2018). Quorum-sensing regulator RhlR but not its autoinducer RhlI enables Pseudomonas to evade opsonization. EMBO Rep 19, e44880.

Haller, S., Limmer, S., and Ferrandon, D. (2014). Assessing Pseudomonas virulence with a nonmammalian host: Drosophila melanogaster. Methods Mol Biol 1149, 723–740.

Hanson, M.A., and Lemaitre, B. (2023). Antimicrobial peptides do not directly contribute to aging in Drosophila, but improve lifespan by preventing dysbiosis. Dis Model Mech 16, dmm049965.

Hay, B.A., Wolff, T., and Rubin, G.M. (1994). Expression of baculovirus P35 prevents cell death in Drosophila. Development 120, 2121–2129.

Herrera, S.C., and Bach, E.A. (2019). JAK/STAT signaling in stem cells and regeneration: from Drosophila to vertebrates. Development 146, 167643.

Hodgson, J.J., Chen, R.Y., Blissard, G.W., and Buchon, N. (2024). Viral and cellular determinants of polarized trafficking of viral envelope proteins from insect-specific and insect-vectored viruses in insect midgut and salivary gland cells. J Virol 98, e0054024.

Iatsenko, I., Boquete, J.P., and Lemaitre, B. (2018). Microbiota-Derived Lactate Activates Production of Reactive Oxygen Species by the Intestinal NADPH Oxidase Nox and Shortens Drosophila Lifespan. Immunity 49, 929–942 e925.

Jiang, H., and Edgar, B.A. (2011). Intestinal stem cells in the adult Drosophila midgut. Exp Cell Res 317, 2780–2788.

Jiang, H., Patel, P.H., Kohlmaier, A., Grenley, M.O., McEwen, D.G., and Edgar, B.A. (2009). Cytokine/Jak/Stat signaling mediates regeneration and homeostasis in the Drosophila midgut. Cell 137, 1343–1355.

Jones, R.M., Luo, L., Ardita, C.S., Richardson, A.N., Kwon, Y.M., Mercante, J.W., Alam, A., Gates, C.L., Wu, H., Swanson, P.A., et al. (2013). Symbiotic lactobacilli stimulate gut epithelial proliferation via Nox-mediated generation of reactive oxygen species. EMBO J 32, 3017–3028.

Jung, A., Criqui, M.-C., Rutschmann, S., Hoffmann, J.-A., and Ferrandon, D. (2001). A microfluorometer assay to measure the expression of ß-galactosidase and GFP reporter genes in single *Drosophila* flies. Biotechniques 30, 594–601.

Kim, S.H., and Lee, W.J. (2014). Role of DUOX in gut inflammation: lessons from Drosophila model of gut-microbiota interactions. Front Cell Infect Microbiol 3, 116.

Lee, K.Z., Lestradet, M., Socha, C., Schirmeier, S., Schmitz, A., Spenle, C., Lefebvre, O., Keime, C., Yamba, W.M., Bou Aoun, R., et al. (2016). Enterocyte Purge and Rapid Recovery Is a Resilience Reaction of the Gut Epithelium to Pore-Forming Toxin Attack. Cell Host Microbe 20, 716–730.

Lemaitre, B., and Miguel-Aliaga, I. (2013). The digestive tract of Drosophila melanogaster. Annu Rev Genet 47, 377–404.

Lemaitre, B., Reichhart, J.M., and Hoffmann, J.A. (1997). Drosophila host defense: differential induction of antimicrobial peptide genes after infection by various classes of microorganisms. Proc Natl Acad Sci U S A 94, 14614–14619.

Lestradet, M., Lee, K.Z., and Ferrandon, D. (2014). Drosophila as a model for intestinal infections. Methods Mol Biol 1197, 11–40.

Lian, S., Liu, J., Wu, Y., Xia, P., and Zhu, G. (2022). Bacterial and Viral Co-Infection in the Intestine: Competition Scenario and Their Effect on Host Immunity. International journal of molecular sciences 23, 2311.

Limmer, S., Haller, S., Drenkard, E., Lee, J., Yu, S., Kocks, C., Ausubel, F.M., and Ferrandon, D. (2011). Pseudomonas aeruginosa RhlR is required to neutralize the cellular immune response in a Drosophila melanogaster oral infection model. Proc Natl Acad Sci U S A 108, 17378–17383.

Liu, Z., Zhang, H., Lemaitre, B., and Li, X. (2024). Duox activation in Drosophila Malpighian tubules stimulates intestinal epithelial renewal through a countercurrent flow. Cell Rep 43, 114109.

Magwire, M.M., Fabian, D.K., Schweyen, H., Cao, C., Longdon, B., Bayer, F., and Jiggins, F.M. (2012). Genome-wide association studies reveal a simple genetic basis of resistance to naturally coevolving viruses in Drosophila melanogaster. PLoS Genet 8, e1003057.

Maillard, P.V., Ciaudo, C., Marchais, A., Li, Y., Jay, F., Ding, S.W., and Voinnet, O. (2013). Antiviral RNA interference in mammalian cells. Science 342, 235–238.

Maillard, P.V., Van der Veen, A.G., Deddouche-Grass, S., Rogers, N.C., Merits, A., and Reis e Sousa, C. (2016). Inactivation of the type I interferon pathway reveals long double-stranded RNA-mediated RNA interference in mammalian cells. EMBO J 35, 2505–2518.

Medzhitov, R., Schneider, D.S., and Soares, M.P. (2012). Disease tolerance as a defense strategy. Science 335, 936–941.

Micchelli, C.A., and Perrimon, N. (2006). Evidence that stem cells reside in the adult Drosophila midgut epithelium. Nature 439, 475–479.

Mondotte, J.A., Gausson, V., Frangeul, L., Blanc, H., Lambrechts, L., and Saleh, M.C. (2018). Immune priming and clearance of orally acquired RNA viruses in Drosophila. Nat Microbiol 3, 1394–1403.

Nagai, H., Nagai, L.A.E., Tasaki, S., Nakato, R., Umetsu, D., Kuranaga, E., Miura, M., and Nakajima, Y. (2023). Nutrient-driven dedifferentiation of enteroendocrine cells promotes adaptive intestinal growth in Drosophila. Dev Cell 58, 1764–1781 e1710.

Nehme, N.T., Liegeois, S., Kele, B., Giammarinaro, P., Pradel, E., Hoffmann, J.A., Ewbank, J.J., and Ferrandon, D. (2007). A model of bacterial intestinal infections in Drosophila melanogaster. PLoS Pathog 3, e173.

Niehus, S., Giammarinaro, P., Liegeois, S., Quintin, J., and Ferrandon, D. (2012). Fly culture collapse disorder: detection, prophylaxis and eradication of the microsporidian parasite Tubulinosema ratisbonensis infecting Drosophila melanogaster. Fly (Austin) 6, 193–204.

Nigg, J.C., Castello-Sanjuan, M., Blanc, H., Frangeul, L., Mongelli, V., Godron, X., Bardin, A.J., and Saleh, M.C. (2024). Viral infection disrupts intestinal homeostasis via Sting-dependent NF-kappaB signaling in Drosophila. Curr Biol 34, 2785–2800 e2787.

Nigg, J.C., Mongelli, V., Blanc, H., and Saleh, M.C. (2022). Innovative Toolbox for the Quantification of Drosophila C Virus, Drosophila A Virus, and Nora Virus. J Mol Biol 434, 167308.

Ohlstein, B., and Spradling, A. (2006). The adult Drosophila posterior midgut is maintained by pluripotent stem cells. Nature 439, 470–474.

Patel, P.H., Penalva, C., Kardorff, M., Roca, M., Pavlovic, B., Thiel, A., Teleman, A.A., and Edgar, B.A. (2019). Damage sensing by a Nox-Ask1-MKK3-p38 signaling pathway mediates regeneration in the adult Drosophila midgut. Nature communications 10, 4365.

Poirier, E.Z., Buck, M.D., Chakravarty, P., Carvalho, J., Frederico, B., Cardoso, A., Healy, L., Ulferts, R., Beale, R., and Reis e Sousa, C. (2021). An isoform of Dicer protects mammalian stem cells against multiple RNA viruses. Science 373, 231–236.

Rahmatika, D., Kuroda, N., Min, Z., Nainu, F., Nagaosa, K., and Nakanishi, Y. (2019). Inhibitory effects of viral infection on cancer development. Virology 528, 48–53.

Rahme, L.G., Stevens, E.J., Wolfort, S.F., Shao, J., Tompkins, R.G., and Ausubel, F.M. (1995). Common virulence factors for bacterial pathogenicity in plants and animals. Science 268, 1899–1902.

Rera, M., Bahadorani, S., Cho, J., Koehler, C.L., Ulgherait, M., Hur, J.H., Ansari, W.S., Lo, T., Jr., Jones, D.L., and Walker, D.W. (2011). Modulation of longevity and tissue homeostasis by the Drosophila PGC-1 homolog. Cell Metab 14, 623–634.

Rogers, A., Towery, L., McCown, A., and Carlson, K.A. (2020). Impaired Geotaxis as a Novel Phenotype of Nora Virus Infection of Drosophila melanogaster. Scientifica (Cairo) 2020, 1804510.

Rutschmann, S., Jung, A.C., Hetru, C., Reichhart, J.M., Hoffmann, J.A., and Ferrandon, D. (2000). The Rel protein DIF mediates the antifungal but not the antibacterial host defense in Drosophila. Immunity 12, 569–580.

Sansone, C.L., Cohen, J., Yasunaga, A., Xu, J., Osborn, G., Subramanian, H., Gold, B., Buchon, N., and Cherry, S. (2015). Microbiota-Dependent Priming of Antiviral Intestinal Immunity in Drosophila. Cell Host Microbe 18, 571–581.

Schissel, M., Best, R., Liesemeyer, S., Tan, Y.D., Carlson, D.J., Shaffer, J.J., Avuthu, N., Guda, C., and Carlson, K.A. (2021). Effect of Nora virus infection on native gut bacterial communities of Drosophila melanogaster. AIMS Microbiol 7, 216–237.

Schneider, J., and Imler, J.L. (2021). Sensing and signalling viral infection in drosophila. Dev Comp Immunol 117, 103985.

Segrist, E., Dittmar, M., Gold, B., and Cherry, S. (2021). Orally acquired cyclic dinucleotides drive dSTING-dependent antiviral immunity in enterocytes. Cell reports 37, 110150.

Segrist, E., Miller, S., Gold, B., Li, Y., and Cherry, S. (2024). Tissue specific innate immune responses impact viral infection in Drosophila. PLoS Pathog 20, e1012672.

Sina Rahme, B., Lestradet, M., Di Venanzio, G., Ayyaz, A., Yamba, M.W., Lazzaro, M., Liegeois, S., Garcia Vescovi, E., and Ferrandon, D. (2022). The fliR gene contributes to the virulence of S. marcescens in a Drosophila intestinal infection model. Scientific reports 12, 3068.

Socha, C., Pais, I.S., Lee, K.Z., Liu, J., Liegeois, S., Lestradet, M., and Ferrandon, D. (2023). Fast drosophila enterocyte regrowth after infection involves a reverse metabolic flux driven by an amino acid transporter. iScience 26, 107490.

Takeishi, A., Kuranaga, E., Tonoki, A., Misaki, K., Yonemura, S., Kanuka, H., and Miura, M. (2013). Homeostatic epithelial renewal in the gut is required for dampening a fatal systemic wound response in Drosophila. Cell Rep 3, 919–930.

Tauszig-Delamasure, S., Bilak, H., Capovilla, M., Hoffmann, J.A., and Imler, J.L. (2002). Drosophila MyD88 is required for the response to fungal and Gram-positive bacterial infections. Nat Immunol 3, 91–97.

Thibault, S.T., Singer, M.A., Miyazaki, W.Y., Milash, B., Dompe, N.A., Singh, C.M., Buchholz, R., Demsky, M., Fawcett, R., Francis-Lang, H.L., et al. (2004). A complementary transposon tool kit for Drosophila melanogaster using P and piggyBac. Nat Genet 36, 283–287.

van Mierlo, J.T., Bronkhorst, A.W., Overheul, G.J., Sadanandan, S.A., Ekstrom, J.O., Heestermans, M., Hultmark, D., Antoniewski, C., and van Rij, R.P. (2012). Convergent evolution of argonaute-2 slicer antagonism in two distinct insect RNA viruses. PLoS Pathog 8, e1002872.

van Rij, R.P., Saleh, M.C., Berry, B., Foo, C., Houk, A., Antoniewski, C., and Andino, R. (2006). The RNA silencing endonuclease Argonaute 2 mediates specific antiviral immunity in Drosophila melanogaster. Genes Dev 20, 2985–2995.

Webster, C.L., Waldron, F.M., Robertson, S., Crowson, D., Ferrari, G., Quintana, J.F., Brouqui, J.M., Bayne, E.H., Longdon, B., Buck, A.H., et al. (2015). The Discovery, Distribution, and Evolution of Viruses Associated with Drosophila melanogaster. PLoS Biol 13, e1002210.

Wu, Q., Luo, Y., Lu, R., Lau, N., Lai, E.C., Li, W.X., and Ding, S.W. (2010). Virus discovery by deep sequencing and assembly of virus-derived small silencing RNAs. Proc Natl Acad Sci U S A 107, 1606–1611.

Wu, X., Dao Thi, V.L., Huang, Y., Billerbeck, E., Saha, D., Hoffmann, H.H., Wang, Y., Silva, L.A.V., Sarbanes, S., Sun, T., et al. (2018). Intrinsic Immunity Shapes Viral Resistance of Stem Cells. Cell 172, 423–438 e425.

Wu, X., Kwong, A.C., and Rice, C.M. (2019). Antiviral resistance of stem cells. Curr Opin Immunol 56, 50–59.

Xu, R., Lou, Y., Tidu, A., Bulet, P., Heinekamp, T., Martin, F., Brakhage, A., Li, Z., Liegeois, S., and Ferrandon, D. (2023). The Toll pathway mediates Drosophila resilience to Aspergillus mycotoxins through specific Bomanins. EMBO Rep 24, e56036.

Zeng, X., Chauhan, C., and Hou, S.X. (2010). Characterization of midgut stem cell- and enteroblast-specific Gal4 lines in drosophila. Genesis 48, 607–611.

Zhou, F., Rasmussen, A., Lee, S., and Agaisse, H. (2013). The UPD3 cytokine couples environmental challenge and intestinal stem cell division through modulation of JAK/STAT signaling in the stem cell microenvironment. Dev Biol 373, 383–393.

