## Supplementary Legends for "Nora virus proliferates in dividing intestinal stem cells and thereby sensitizes *Drosophila* flies to *Pseudomonas aeruginosa* intestinal infection and to oxidative stress"

### Supplementary Figures Legends

**Supplementary Figure 1. *pastrel* is not involved in the *P. aeruginosa* infection enhanced sensitivity of Nora virus-infected flies**

(**A**) Survival analysis of Ore-R(SM) (Nora(+)) and Ore-R(SC) (Nora(-)) flies following intestinal infection with *S. marcescens* Db11 at 25 °C compared with non-infected control flies (NI) maintained on a sucrose-only diet. (**B**) Survival analysis of Ore-R(SM) (Nora(+)) and Ore-R(SC) (Nora(-)) flies maintained on a sucrose-only diet at 25 °C. (**C**) Nora virus RNA levels measured by RT-qPCR in multiple wild-type fly lines (Oregon-R (SC), *ywDD1*, *cn bw*, Canton-S) naturally infected with Nora virus. (**D**) PCR analysis of *pastrel* (*pst*) alleles associated with sensitivity ($pst^S$) or resistance ($pst^R$) in the wild-type stocks analyzed in (A-C). Primer sequences and quantification strategy are described in the Methods section. (**E-H**) Survival analysis following *P. aeruginosa* PA14 intestinal infection at 25 °C in Nora-infected flies from the indicated genetic backgrounds: w[A5001] (E), Canton-S (F), *ywDD1*, *cn bw* (G), and Oregon-R(SM) (H). (**I**) Survival analysis following PA14 intestinal infection at 25 °C in Nora-infected Oregon-R(SC) flies. Each graphic is representative of three independent experiments. Survival panels are presented as mean ± SEM with each curve representing a triplicate of 20 flies. Other panel is presented as box and whiskers where the middle bar of the box plots represents the median, and the upper and lower limits of boxes indicate the first and third quartiles, respectively; the whiskers define the minima and maxima. Each dot in Nora virus load panel (C) represents a sample of 5 flies. Survival data were analyzed using a log-rank (Mantel-Cox) test. Nora virus load quantification (C) was analyzed using one-way ANOVA with Tukey's post-hoc test. Statistical significance is indicated as ***p < 0.001, ****p < 0.0001.

**Supplementary Figure 2. Infection of Nora(-) flies with Nora virus through fecal contamination recapitulates the fitness properties of Nora naturally infected flies**

Flies used in these experiments were derived from stocks cured of Nora virus by egg bleaching. A subset of cured flies was re-exposed to Nora virus via fecal material collected from the original Nora-infected Ore-R stock (Nora(+) re-inf feces), while control flies remained uninfected (Nora(-)). *P. aeruginosa*: PA14; non-infected controls: NI.

(**A**) Nora virus RNA levels measured by RT-qPCR in Nora(+) and Nora(-) flies across successive generations (G0, G1, and G2) following fecal-mediated re-infection. (**B**) Survival analysis of re-infected Nora(+) and Nora(-) flies following oral exposure to PA14 at 25 °C. (**C**) Quantification of phospho-histone H3 (PHH3)-positive nuclei in the midgut of re-infected Nora(+) and Nora(-) flies at 2 days following PA14 intestinal infection or in non-infected controls. Each graphic is representative of three independent experiments. Survival panel is presented as mean ± SEM with each curve representing a triplicate of 20 flies. Other panels are presented as box and whiskers where the middle bar of the box plots represents the median, and the upper and lower limits of boxes indicate the first and third quartiles, respectively; the whiskers define the minima and maxima. Each dot in Nora virus load panel (A) represents a sample of 5 flies. Each dot in the PHH3 quantification panel (C) represents one single posterior midgut. Survival data (B) were analyzed using a log-rank (Mantel-Cox) test. Nora virus load (A) and PHH3 quantifications (C) were analyzed using one-way ANOVA with Tukey's post-hoc test. Statistical significance is indicated as **p < 0.01, ***p < 0.001, ****p < 0.0001.

**Supplementary Figure 3. Influence of age, diet, and microbiota on Nora virus load, survival to *P. aeruginosa* infection, and fly fitness**

(**A-B**) Survival analysis of Nora-infected (Nora(+)) and non-infected (Nora(-)) flies following *P. aeruginosa* PA14 intestinal infection. Flies were raised either on standard food (A) or were raised on rich food (standard medium supplemented with 5x yeast) (B); they were transferred to a sucrose-only diet just prior to infection. (**C**) Quantification of PA14 colony-forming units (CFUs) in whole flies raised on standard or rich food, comparing Nora(+) and Nora(-) flies. (**D**) Quantification of phospho-histone H3 (PHH3)-positive nuclei in the posterior midgut at 3 days post-PA14 infection from experiments shown in (A) and (B). (**E**) Comparison of PHH3-positive nuclei between young (5-day-old) and aged (30-day-old) Nora(+) and Nora(-) flies. (**F**) Quantification of microbial load in midguts of Nora(+) and Nora(-) flies maintained on a sucrose-only diet for 7 days. (**G**) Survival analysis of Nora(-) and Nora(+) flies maintained on sucrose diet with or without antibiotic treatment (ABX). (**H**) Quantification of PHH3-positive nuclei in midguts of Nora(+) and Nora(-) flies maintained on a sucrose-only diet for 7 days, in the presence or absence of antibiotics. Each graphic is representative of three independent experiments. Survival panels are presented as mean ± SEM with each curve representing a triplicate of 20 flies. Other panels are presented as box and whiskers where the middle bar of the box plots represents the median, and the upper and lower limits of boxes indicate the first and third quartiles, respectively; the whiskers define the minima and maxima. Each dot in PHH3 quantification panels (D-E, H) represent one single posterior midgut. Each dot in the PA14 CFU quantification panel (C) represents one single posterior midgut. Each dot in the CFU quantification panel (F) represents one single posterior midgut. Survival data (A-B, G) were analyzed using a log-rank (Mantel-Cox) test. PA14 CFU counts (C) and PHH3 quantifications (D-E) were analyzed using one-way ANOVA with Tukey's post-hoc test.

CFU counts (F) were analyzed using the Mann-Whitney nonparametric test. Statistical significance is indicated as $^{**}p < 0.01$, $^{***}p < 0.001$, $^{****}p < 0.0001$.

**Supplementary Figure 4. Effects of Nora virus on survival and epithelial proliferation following various systemic infections or sterile injury**

(**A**) Survival analysis of Nora-infected (Nora(+)) and non-infected (Nora(-)) flies following natural infection with the fungus *Beauveria bassiana*. *MyD88* mutant flies: Toll mutant positive control; w[A5001] flies: wild-type control. All control and Nora(-) experimental flies were confirmed negative for Nora virus. (**B**) Survival analysis of Nora(+) and Nora(-) flies following septic injury with *Enterococcus faecalis*. (**C**) Survival analysis of Nora(+) and Nora(-) flies following septic injury with *Enterobacter cloacae*. *key* flies: IMD pathway mutant positive control; w[A5001] flies: wild-type control. (**D**) Survival of Nora(+) and Nora(-) flies following sterile PBS injection at 29 °C. (**E**) Quantification of PHH3-positive nuclei in midguts of Nora(+) and Nora(-) flies at 4 days following sterile PBS injection (clean injury, CI) or without injury (NI) at 29 °C. (**F**) Survival to clean injury of Nora(+) and Nora(-) flies. Each graphic is representative of three independent experiments. Survival panels are presented as mean ± SEM with each curve representing a triplicate of 20 flies. Other panel is presented as box and whiskers where the middle bar of the box plots represents the median, and the upper and lower limits of boxes indicate the first and third quartiles, respectively; the whiskers define the minima and maxima. Each dot in the PHH3 quantification panel (E) represents one single posterior midgut. Survival data (A-D, F) were analyzed using a log-rank (Mantel-Cox) test. PHH3 quantification (E) was analyzed using one-way ANOVA with Tukey's post-hoc test. Statistical significance is indicated as $^{**}p < 0.01$, $^{****}p < 0.0001$.

**Supplementary Figure 5. Effects of age, diet, and stress on Nora virus abundance in the intestine**

(**A**) Representative confocal images of 5-day-old non-infected fly intestines. Tissues were fixed and stained for DNA (DAPI, blue), actin (FITC, green), and Dicer-2 (mRFP, red). Scale bar, 5 µm. Note the nonspecific staining in the visceral muscles due to the secondary antibody. (**B**) Representative confocal image from 5-day-old Nora-infected (+) fly intestines, showing either one enteroblast or an enteroendocrine cell. Scale bar, 5 µm. (**C**) Representative confocal image of non-infected mRFP::Dicer-2 flies intestines. The progenitor cells are outlined. Tissues were stained for DNA (DAPI, blue), actin (FITC, green), and Dicer-2 (mRFP, red). Scale bar, 5 µm. (**D**) Quantification of Nora-positive cells in the intestine after infectious (*P. aeruginosa* PA14, *S. marcescens* Db11) or chemical (1 mM paraquat) stresses. (**E**) Representative confocal images of 30-day-old Nora-infected fly intestines. Scale bars, 3 µm (left) and 5 µm (right). (**F**) Quantification of Nora-positive cells in intestines from young and aged flies. (**G**) Quantification of Nora(+) cell types of the intestinal epithelium according to age. EC: enterocyte; ISC: progenitor cell (**H**) Representative confocal images of 5-day-old Nora-infected intestines of flies maintained on standard food (left) or rich food and PA14 infection (right). Scale bar, 5 µm. (**I**) Quantification of Nora-positive cells shown in (E & G). (**J**) Scheme of the *Drosophila* fore- and midgut displaying the R4&R5 regions in which Nora-positive cells have been detected. Each graphic is representative of three independent experiments. Panels are presented as box and whiskers where the middle bar of the box plots represents the median, and the upper and lower limits of boxes indicate the first and third quartiles, respectively; the whiskers define the minima and maxima. Each dot in the Nora virus quantification panels (D, F, H) represent a region of one single posterior midgut. Nora-positive cell quantifications were analyzed using Mann-Whitney nonparametric test (F, I) or one-way ANOVA with Tukey's post-hoc test (D, G). Statistical significance is indicated as $^{**}p < 0.01$, $^{****}p < 0.0001$.

**Supplementary Figure 6. Effects of Nora virus and stress conditions on intestinal cell death**

(**A-C**) Representative confocal images of midguts from Nora-infected (Nora(+)) flies, labeled with DAPI (blue) and ApopTag to detect apoptotic cells. Images show representative midguts under different conditions: without stimulation (A), following *P. aeruginosa* PA14 intestinal infection (B), and following 1 mM paraquat exposure (C). Cell death quantification corresponding to these conditions is presented in Figure 6A.
