## Supplementary figures and images for "Nora virus proliferates in dividing intestinal stem cells and thereby sensitizes *Drosophila* flies to *Pseudomonas aeruginosa* intestinal infection and to oxidative stress"

### Supplementary Figures

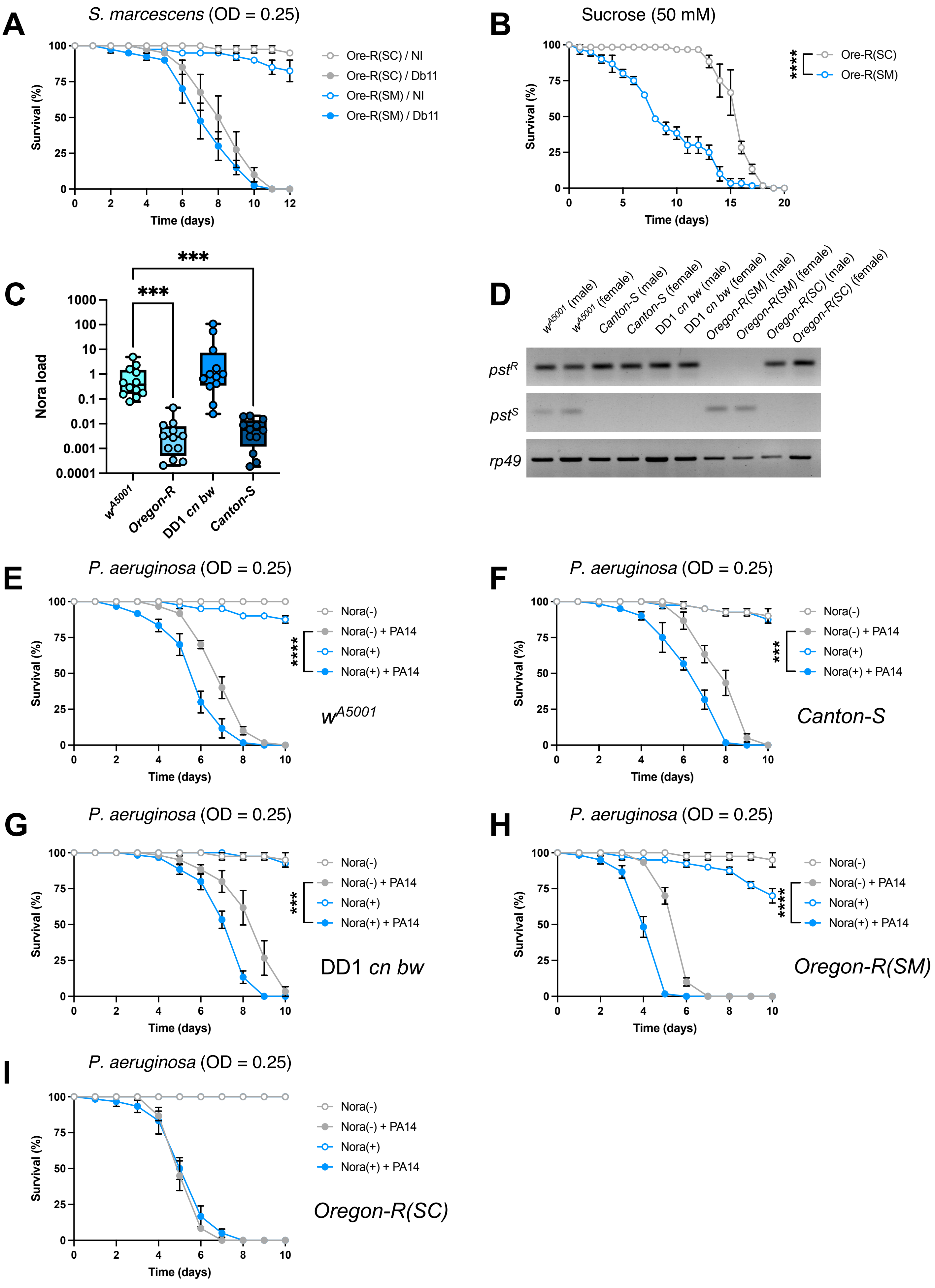

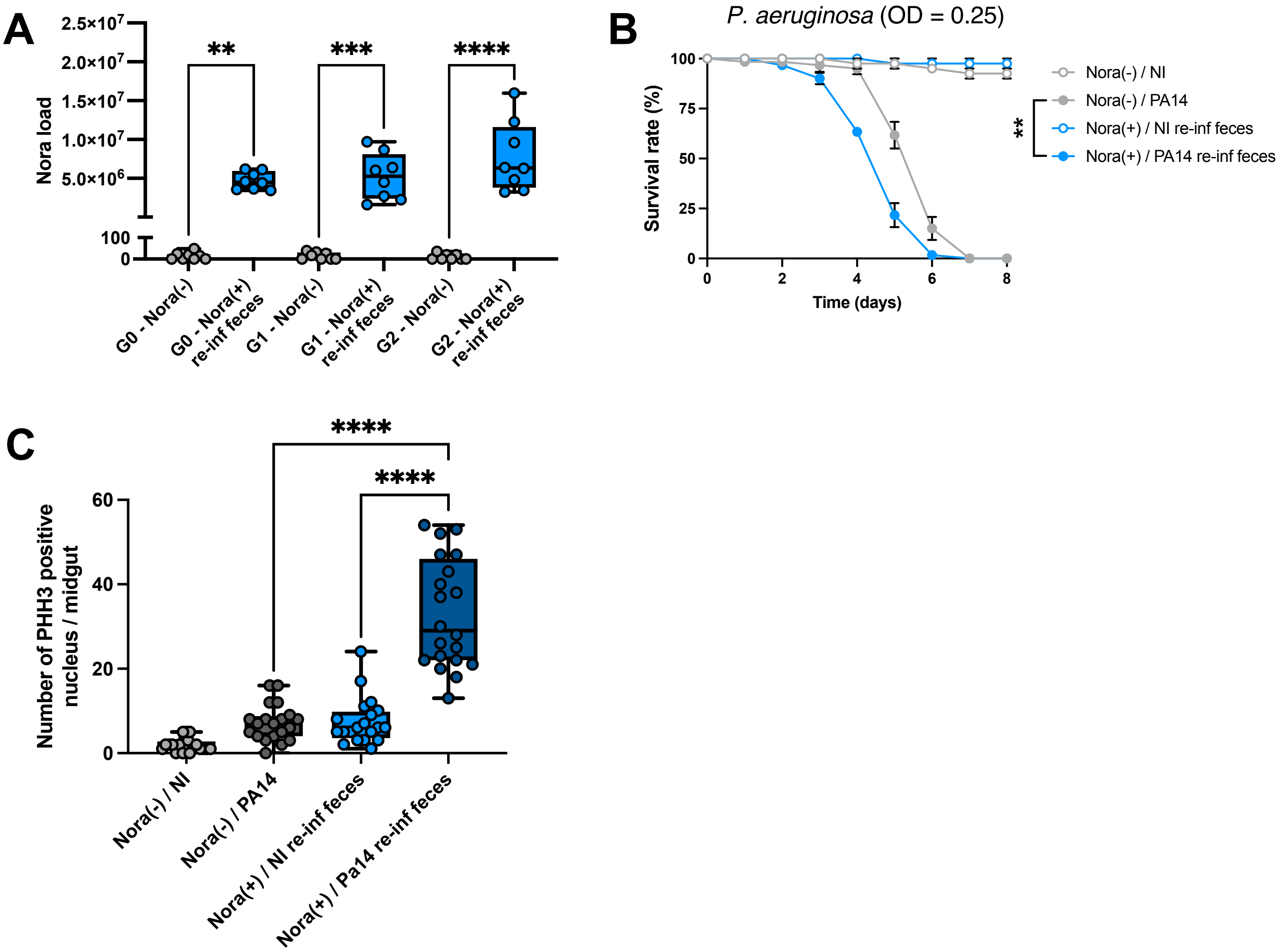

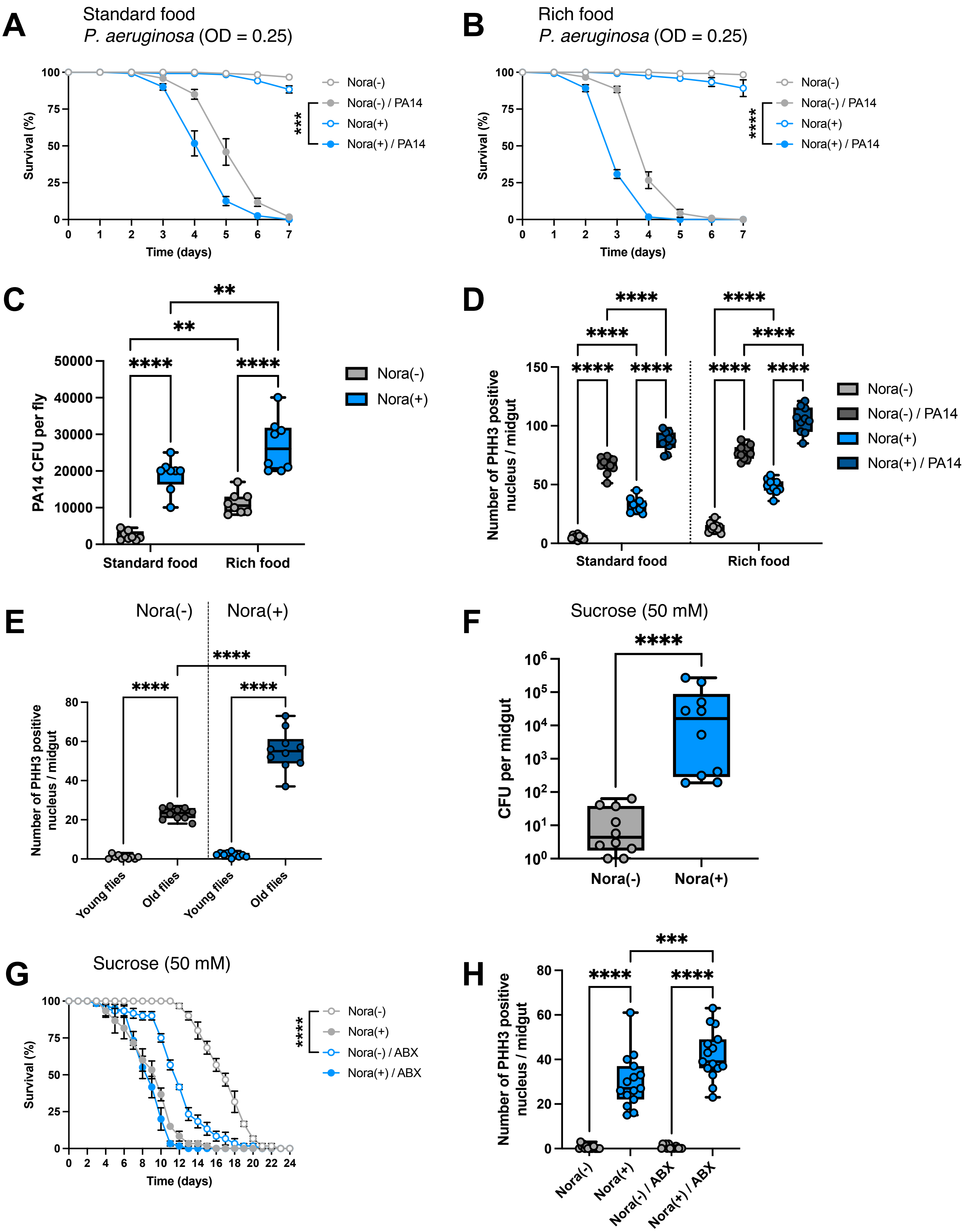

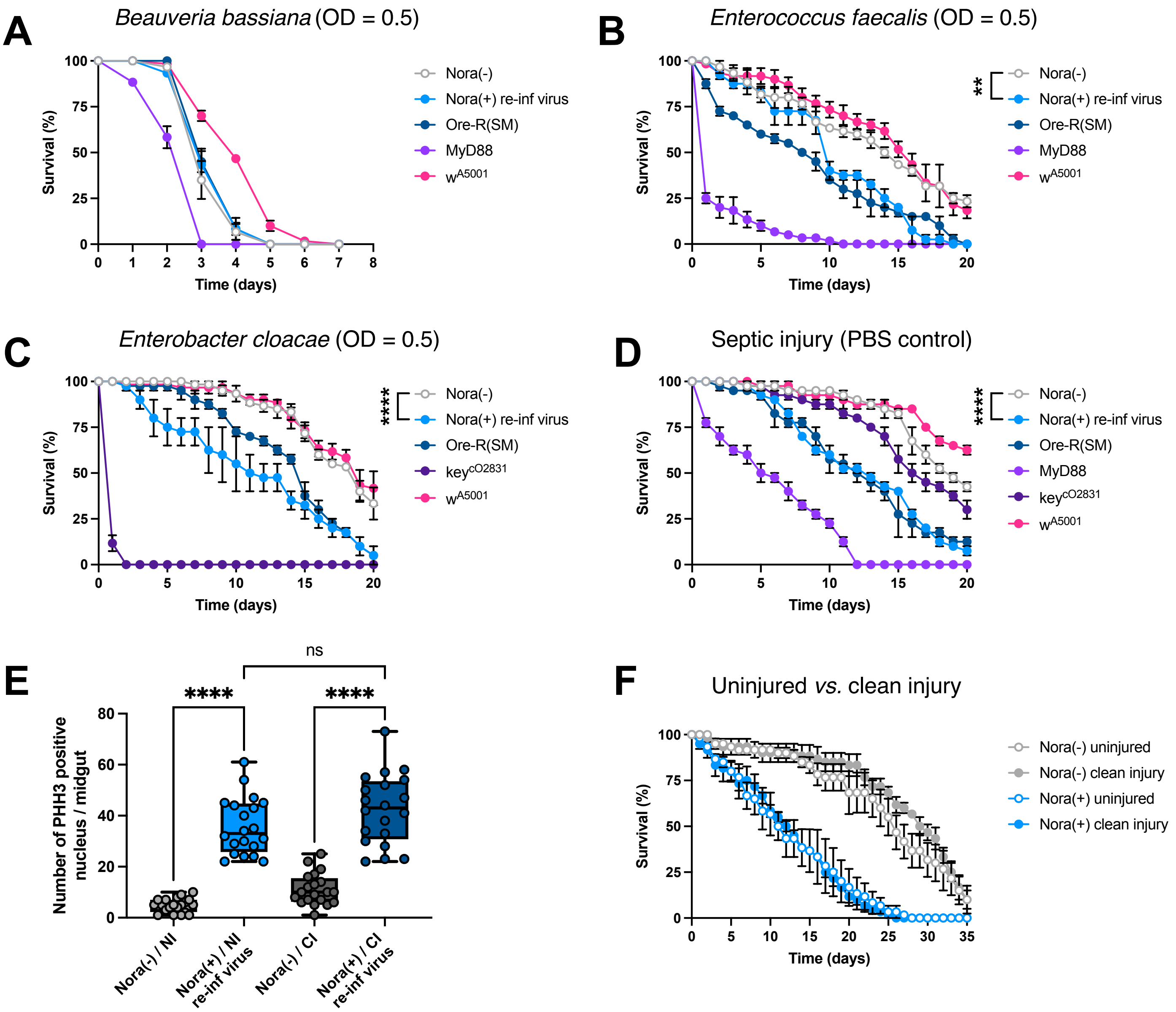

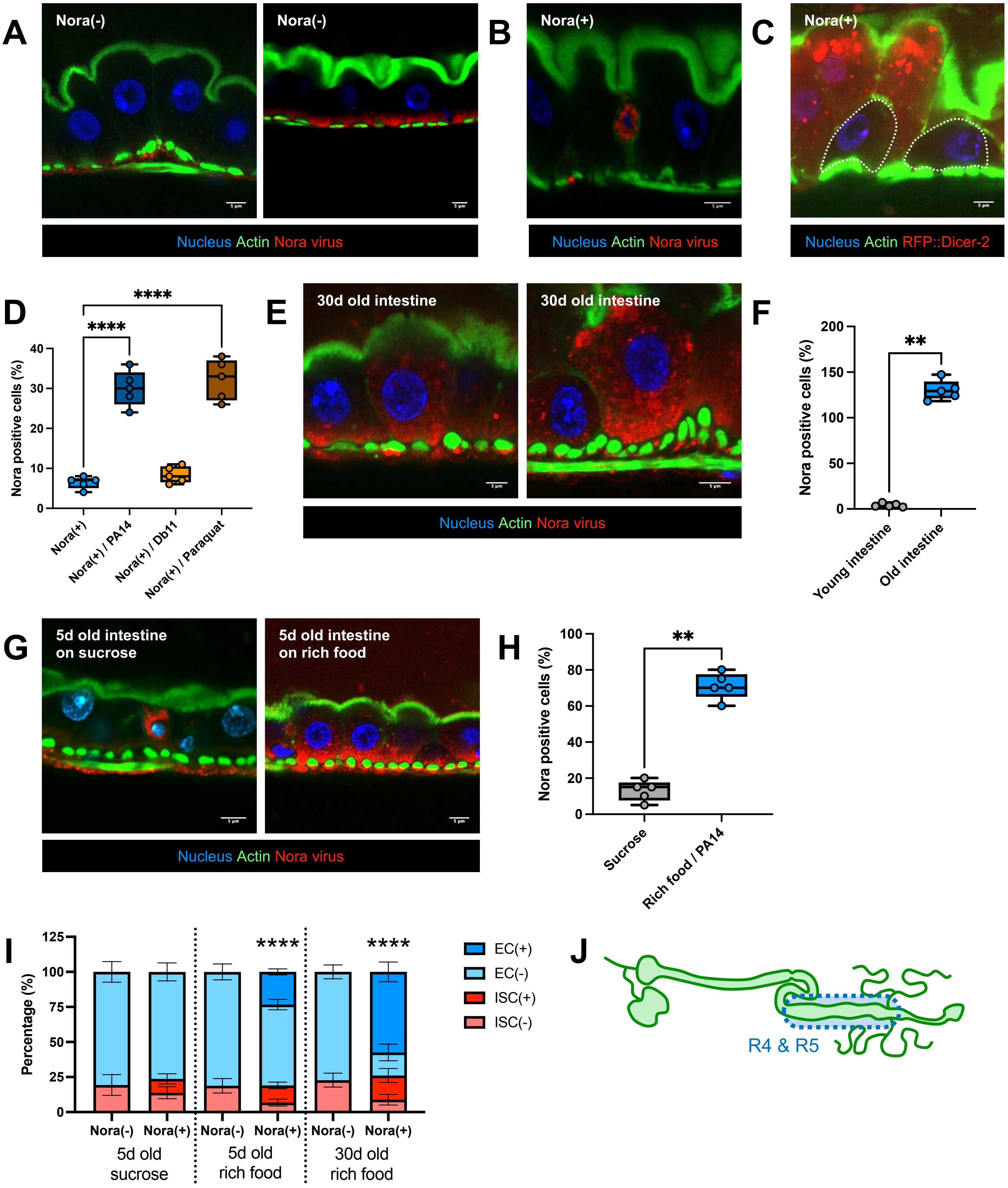

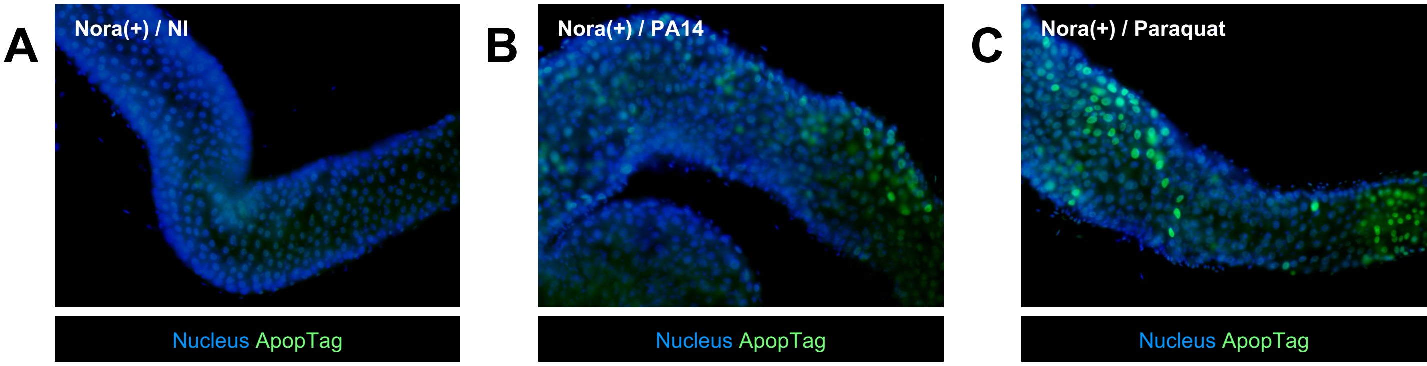
