## Supplementary material for "Nora virus proliferates in dividing intestinal stem cells and thereby sensitizes *Drosophila* flies to *Pseudomonas aeruginosa* intestinal infection and to oxidative stress": Table

| Name | Sequence (5’-3’) |
|---|---|
| rp49 Fw (also known as RpL32) | GACGCTTCAAGGGACAGTATCTG |
| rp49 Rv (also known as RpL32) | AAACGCGGTTCTGCATGAG |
| Nora Fw | AACCTCGTAGCAATCCTCTCAAG |
| Nora Rv | TTCTTGTCCGGTGTATCCTGTATC |
| Diptericin Fw | GCTGCGCAATCGCTTCTACT |
| Diptericin Rv | TGGTGGAGTGGGCTTCATG |
| upd3 Fw | CGACCTGCAGATTTACGTGG |
| upd3 Rv | GGTCCCAGTGCAACTTGATG |
| dome Fw | TGACCGATACATTCCGCGTA |
| dome Rv | GTAGGTGATGGGCTTCTCGT |
| hop Fw | GATGAGACCAAGCGCTTCAG |
| hop Rv | TTTCCGCCTTGATCGTGTTG |
| Stat92E Fw | CGTTACGCGCAATACACAGA |
| Stat92E Rv | CGTTTTGAATCTCGCCCGAT |
| Socs36E Fw | GACAAATCGAACTGCTGGCA |
| Socs36E Rv | CGTTGTTATTCACGGGCTGT |
| PIAS Fw (also known as Su(var)2-10) | ACTGTCTGGCCGTATACCTG |
| PIAS Rv (also known as Su(var)2-10) | CCCTTCGTCTTCATTCGCTG |
| DCV Fw | TCATCGGTATGCACATTGCT |
| DCV Rv | CGCATAACCATGCTCTTCTG |
| FHV.1 Fw | TTTAGAGCACATGCGTCCAG |
| FHV.1 Rv | CGCTCACTTTCTTCGGGTTA |
| FHV.2 Fw | CAACGTCGAACTTGATGCAG |
| FHV.2 Rv | GCTTTACAGGGCATTTCCAA |
| VSV Fw | CATGATCCTGCTCTTCGTCA |
| VSV Rv | TGCAAGCCCGGTATCTTATC |
| Sinv Fw | CAAATGTGCCACAGATACCG |
| Sinv Rv | ATACCCTGCCCTTTCAACAA |
| CrPV-1 Fw | GCTGAAACGTTCAACGCATA |
| CrPV-1 Rv | CCACTTGCTCCATTTGGTTT |
| CrPV-2 Fw | GGAATTTTTGGAGACGCAAA |
| CrPV-2 Rv | GTGAAGGGGGCAACTACAAA |
| DAV-1 Fw | CGAACTGCCAACTGAGGTCT |
| DAV-1 Rv | CCACCCCGGTTGTTAATGGA |
| IIV6 Fw | TTGTTAGGAATTGGAACTGGAA |
| IIV6 Rv | GCCCTAGATGCTGCTTGTTC |
| DBV-A Fw | TGCAGTCAGACGCCAGTATC |
| DBV-A Rv | CCCCTGAACCTGGTAGCATA |
| DTV Fw | AGTTTTGGGATTGGCAACAG |
| DTV Rv | TTACGCCCCTTGAATGGTAG |
| DXV-A Fw | CATCGTCGACATCACCAAAC |
| DXV-A Rv | TCCTGTGAAAGCTGCAAATG |
| DXV-B Fw | TGAGCAAAATTACGCACAGG |
| DXV-B Rv | CCATACGCGTTGTGTATTCG |
